# The global explosion of eukaryotic algae: the potential role of phosphorus?

**DOI:** 10.1101/2020.05.27.118547

**Authors:** Lisa K Eckford-Soper, Donald E. Canfield

**Affiliations:** Nordcee, Department of Biology, University of Southern Denmark, Odense M, Denmark

## Abstract

There arose one of the most important ecological transitions in Earth’s history approximately 750 million years ago during the middle Neoproterozoic Era (1000 to 541 million years ago, Ma). Biomarker evidence suggests that around this time there was a rapid shift from a predominantly bacterial-dominated world to more complex ecosystems governed by eukaryotic primary productivity. The resulting ‘Rise of the algae’ led to dramatically altered food webs that were much more efficient in terms of nutrient and energy transfer. Yet, what triggered this ecological shift? In this study we examined the theory that it was the alleviation of phosphorus (P) deficiency that gave eukaryotic alga the prime opportunity to flourish. We performed laboratory experiments on the cyanobacterium *Synochystis salina* and the eukaryotic algae *Tetraselmis suecia* and examined their ability to compete for phosphorus. Both these organisms co-occur in modern European coastal waters and are not known to have any allelopathic capabilities. The strains were cultured in mono and mixed cultures in chemostats across a range of dissolved organic phosphorus (DIP) conditions to reflect modern and ancient oceanic conditions of 0.2 μM P and 2 μM P, respectively. Our results show that the cyanobacteria outcompete the algae at the low input (0.2 μM P) treatment, yet the eukaryotic algae were not completely excluded and remained a constant background component in the mixed-culture experiments. Also, despite it’s relatively large cell size, the algae *T. suecia* had a high affinity for DIP. With DIP input concentrations resembling modern-day levels (2 μM), the eukaryotic algae could effectively compete against the cyanobacteria in terms of total biomass production. These results suggest that the availability of phosphorus could have influenced the global expansion of eukaryotic algae. However, P limitation does not seem to explain the complete absence of eukaryotic algae in the biomarker record before ca. 750 Ma.

## Introduction

Molecular clock data indicates that the *Archaeplastida*, the major group of autotrophic eukaryotes comprising of the red algae, the green algae and the common ancestor of all protists appeared somewhere around 1900 Ma, while crown group *Rhodophyta* evolved sometime between 1,600–1,000 Ma [1]. Despite this, the ratio of steranes to hopanes in ancient sediments suggests the eukaryotic algae failed to make any significant biological or ecological impact until well into the Neoproterozoic Era, ca. 750 Ma [2,3]. The Neoproterozoic Era was characterised by extreme biochemical and climatic volatility, which resulted in dramatic alterations in the marine redox state and fluctuating surface-ocean oxygen concentrations [4]. Indeed, during the Neoproterozoic Era and triggered by three major interconnected events, the Earth experienced some of the greatest biological and geochemical changes in its history [5]. Firstly, several massive glaciation episodes occurred, the so called ‘snowball Earth’ events. These glaciations not only altered the Earth’s climate, but they resulted in extensive continental weathering that may have released large quantities of nutrients into the oceans [2,6–8]. Secondly, in the late Neoproterozoic Era, a shift in ecosystem structure and function resulted in the dramatic expansion and diversification of eukaryotic algae, the so called ‘rise of the algae’ [2,3,9]. This ecological transition resulted in irreversibly altered benthic and pelagic ecosystems and the eventual emergence of metazoan life. Lastly, there was an apparent widespread oxygenation of the Earth’s surface environment [10–14].

There are several suggestions as to what triggered the ecological shift leading to eukaryote-dominated productivity. These suggestions range from oxygen concentrations inhibitory to eukaryotes [15], to an increase in predation pressure from the evolution of protist predators [16], and as noted above, an increase in P availability [17]. Phosphorus, unlike fixed nitrogen (nitrate, nitrite, ammonium), cannot be produced biologically [18], and the main source is from the weathering of continental rocks. Due to its essential role in governing protein synthesis, nucleic acid production, adenosine phosphate transformations and intracellular transport, phosphorus may have been the main limiting nutrient controlling primary production through much of Earth’s history since the first rise of atmospheric oxygen known as the Great Oxidation Event (GOE) [19,20].

It has been argued that during the Mesoproterozoic Era (1600 to 1000 Ma), phosphorus scavenging by ferrous iron in anoxic deep waters may have led to the removal of phosphorus from ocean waters, reducing the total phosphorus inventory to concentrations much lower than today [21]. If true, levels of phosphorus may have been low enough to limit primary productivity and thus organic carbon burial, leading to low atmospheric oxygen levels [21–23]. Low phosphate concentrations would have also benefited smaller classes of phytoplankton, including cyanobacteria. With their relatively smaller cell sizes and greater surface area to volume ratios, cyanobacteria would have a physiological advantage, allowing them to outcompete larger eukaryotic algae under these low nutrient conditions [24,25].

During the middle Neoproterozoic Eon (800-650 Ma), fundamental shifts in the phosphorus cycle may have resulted in increased marine P concentrations [7,21,26]. Since, the apparent increase in P concentration occurred around the same time as the first appearance of algal steranes in the biomarker record (780-729 Ma) [3], the two events could be linked [2]. As the availability of phosphorus is regarded as a critical factor regulating phytoplankton and their communities [24], the response of different organisms to nutrient availability should ultimately impact overall community structure [27]. However, a better understanding on how the extent of P-limitation regulates growth and species composition is needed to assess its role in regulating phytoplankton productivity, diversity and succession in the ancient oceans.

In order to effectively exploit a variable P supply, many phytoplankton species have developed an array of mechanisms to cope with low P concentrations. These include: the alteration of cellular P requirements through the substitution of phospholipids with sulphur-based lipids, altered P uptake rates and intracellular P stores, and the utilisation of organic P sources through the release of extracellular enzymes like alkaline phosphatase (AP) [28]. Phytoplankton cells monitor their environment through a feedback system that can simultaneous sense external and internal P concentrations to alter the number and type of cellular P transporters [29] and AP [28].

To gain insight into how phosphorus limitation could have affected phytoplankton population distributions in ancient oceans, we examined the dual hypotheses that: 1) before the rise of algal phosphorus limitation favoured the dominance of cyanobacteria, and 2) the alleviation of phosphorus deficiency triggered the global expansion of eukaryotic algae. Therefore, in this study we examined how cyanobacteria and eukaryotic algae react and adapt to altered phosphorus concentrations by culturing them under a range of P availabilities. Our experimental conditions were chosen to compare and contrast modern ocean conditions with those estimated for the ancient oceans. A green alga and a cyanobacteria were chosen as our model organisms. Both of these organisms have been found to co-occur in modern European coastal waters, both are able to withstand low P environments and neither are known to have any allelopathic capabilities. Experiments were carried out in continuous mixed- and mono-culture experiments.

## Methods

### Cultures

Pure, non-axenic (meaning some bacteria are present) cultures of the alga *Tetraselmis suecia* (CCMP 904) and the cyanobacteria *Synechocystis salina* (CCBA MA001) were obtained from NCMA at Bigelow Laboratory and the culture collection of Baltic algae, respectively. Both species were tested for allelopathic abilities using the methods described in [30]. Allelopathy describes the process where an organism produces chemicals that influence the growth and survival of another [30]. Both strains were grown in a modified BG-11 medium with additional L1 vitamins. Strains were cultured at 15 °C under a light intensity of 80 μmol photons m^−2^ s^−1^ and a 12 h:12 h light:dark cycle. Cultures were grown in 250 mL flasks in batch mode and sequentially acclimated to four different external phosphorus concentrations (100, 5, 2 and 0.2 μM P). Cell numbers were taken daily, and acclimation was complete when cultures exhibited a constant maximum specific growth rate which was calculated during the exponential growth phase.

To calculate the empirical growth rate (μ), cells were cultured until they reached their post-stationary death phase. Each day 1.5 mL of culture was aseptically removed and fixed in 1 % (FC) of acidified Lugol’s iodine and counted via light microscopy [31]. The empirical growth rate was defined as the number of divisions per day^−1^ (Eq. 1). The duration of the exponential growth phase is determined by calculating the maximum achievable R^2^ when fitting straight lines to the logged plots of cell density. The empirical growth rate is defined as:

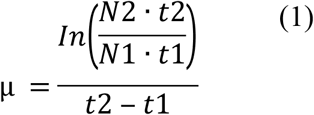

Where Nt_1_ and Nt_2_ are cell abundances at time 1 (t_1_) and time 2 (t_2_), respectively.

### Chemostat Experiments

Cultures were grown in mono and mixed culture in 1 L chemostats. The inflow and outflow rates were controlled by peristaltic pump. Fresh medium was provided at two different phosphorus concentrations of 0.2 μM and 2 μM. The growth rate was set at 0.1 div day^−1^ at 0.2 μM P, and 0.2 div day^−1^ at 2 μM P, equivalent to flow rates of 69 and 138 μL min^−1^ respectively. These rates were set based on growth rates calculated from batch culture (see results). In order to examine the physiological responses of how both species competed for P, they were cultured both separately and together. In the mono-culture experiments *T. suecia* and *S. salina* were grown separately, while in the co-culture experiments, they were grown in the same chemostat.

Inoculums for the experiments were taken from acclimated late-exponential-phase batch stock cultures. As cell sizes differed between the two organisms, they were inoculated into the chemostat with the same total biomass. Bacterial contamination was monitored throughout by staining (DAPI) and with epifluorescent microscopy, following the protocols in [32]. Constant bubbling of filtered air through the chemostats ensured mixing and gas exchange. The pH was measured using a pH meter (Radiometer Analytical, Hach, CO, USA) and maintained at 8 +/− 0.3 throughout. All experiments were carried out in triplicate (n=3).

Every second day, subsamples were removed aseptically from each culture vessel. Aliquots (1.5 mL) were preserved in Lugol’s iodine (1 % FC). Samples containing *T. suecia* were enumerated using a 1 mL Sedgewick Rafter counting chamber, whilst the samples containing *S. salina* were enumerated using a hemocytometer (Burker Turk). In both cases enumeration was performed using a Leica DM 2000 microscope. The mixed samples were therefore counted twice, once for each species. In addition, cell size measurements were also taken to calculate biomass and surface area. For this *T. suecia* was treated as a prolate spheroid [33].

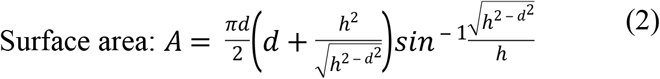

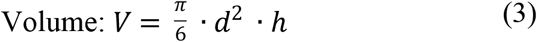

where *d* is the diameter and *h* is the height of *T. suecia*.

*Synochystis salina* can be found in two forms, where approximately 70 % of cells are spherical:

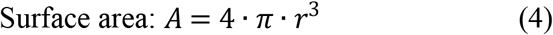

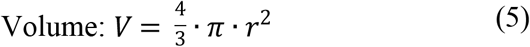

The remaining 30 % are an ellipsoid form with a transapical constriction which can be thought of as a snowman shape. For this, the volume of the two domes were calculated.

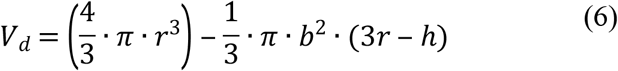

V_d_ being the volume of the dome, *r* is the radius, *b* is the height from the bottom of the sphere to the constriction. The total volume is then given by:

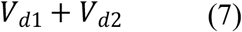

The area is given by:

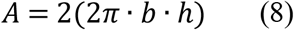

The cell sizes were corrected for shrinkage caused by the fixative. This was done by measuring live cells, immobilised in glycerol, and by comparing these sizes to those measured on Lugol’s fixed cells. The shrinkage caused by Lugol’s iodine was between 11-19 %.

To determine biomass, as well as extracellular and intracellular nutrient and chlorophyll *a* (Chl *a*) concentrations, subsamples (50 mL) of culture were removed aseptically from each reaction vessel and filtered through 25 mm diameter pre-combusted (450 °C for 4 h) GF/F filters [34]. An aliquot (15 mL) of the filtrate was removed for total phosphorus (TP) analysis as described below. A further 3 mL was set aside and kept at ambient temperature (15 °C) for alkaline phosphatase activity (APA) determination. The filtrates and filters for nutrient analysis were frozen at −20 °C for subsequent analysis. Particulate organic carbon (POC) measurements were made as described in [35] using a Thermo Fisher Elemental Analyser and calibrated with isoleucine.

### Chlorophyll *a* Analysis

The filters for Chl *a* analysis were kept in 2 mL Eppendorf® tubes wrapped in aluminium foil. For analysis, the filters were transferred to 15 mL centrifuge tubes, and acetone (8 mL, 90 %) was added to each tube. The samples were kept overnight (5 °C) before sonication (30 min) in a sonication bath and then centrifuged (3000 rpm at 6 °C for 5 min). Chlorophyll *a* was measured with a Turner TD-700 fluorometer (Turner Design, Sunnyvale, CA, USA). The fluorometer was calibrated using a Chl *a* extract from spinach and serial dilutions of a 4 mg L^−1^ stock standard. A solid-state secondary standard (SSS) was measured every ten samples. The SSS insert provides a very stable fluorescent signal and is used when measuring Chl *a* to check for fluorometer stability and sensitivity. The detection limit was 1 μg L^−1^.

### Phosphorus Analysis

Particulate organic phosphorus (POP) was measured in triplicate on frozen filters by the ammonium molybdate method after wet oxidation in acid persulphate (Hansen & Koroleff 1999). Wet oxidation was accomplished by suspending the filters in 10 mL of Milli-Q water in 50 mL Teflon Schott bottles and by adding 1.5 mL acid potassium peroxodisulphate for 90 minutes at 121 ºC in an autoclave. Pre-combusted filters were oxidised along with the samples to account for background P concentrations that were subtracted from the sample values. The samples were then cooled to room temperature. Particulate P was measured as liberated orthophosphate, and its concentration was measured with the standard molybdenum blue technique after sample handling with the following procedure [35,36]. Briefly, ascorbic acid (0.4 mL) and mixed reagent (ammonium heptamolybdate tetrahydrate, sulphuric acid and potassium antimony) (0.2 mL) were added to 10 mL of the sample and mixed [37]. After 10-30 minutes, the absorbance was measured spectrographically using a 10 cm glass cuvette (Thermo Scientific Genesys 1OS UV-VIS, Ma USA). The detection limit was 0.015 μmol L^−1^.

Total phosphorus from the frozen cell-free filtrate (medium) was analysed using the standard molybdenum blue technique as described above after the samples were thawed.

### Alkaline Phosphatase Activity

Alkaline phosphatase activity (APA) was measured using 4-methylumbelliferyl phosphate (MUF-P, Sigma-Aldrich) as a fluorogenic substrate following the protocols in [38]. Briefly, MUF-P (3 μL) was added to 3 mL of the filtrate (100 nM, final concentration). The sample was mixed, after which 1 mL of 50 mM borate buffer (pH 10.8) was added, and the sample was mixed again. Fluorescence was measured on a Turner TD-700 fluorometer (Turner Design, Sunnyvale, CA, USA).

### Nutrient Uptake

Nutrient uptake experiments were carried out on both *T. suecia* and *S. salina*. Cells were harvested from dense exponential-phase batch cultures by gentle centrifugation (3000 *g,* 5 min) and were then resuspended and maintained in phosphate-free media for 48 hours. Cell densities were 8 × 10^4^ cells mL^−1^ for *T. suecia* and 2.7 × 10^4^ cells mL^−1^ for *S. salina.* After the starvation period, the cells were added to 600 mL of phosphorus-replete media (100, 5, 2 and 0.2 μM).

At time intervals of 0, 10, 20, 30, 50, 70, 90 and 120 minutes, 1.5 mL of culture was removed and preserved in Lugol’s iodine (1 % FC) for cell counts as described above. For analysis of particulate organic phosphorus (POP) and total dissolved phosphorus (TP) concentrations, aliquots (50 mL) of culture were removed and filtered through a 25 mm diameter pre-combusted (450 °C, 4 h) GFF filters. An aliquot (15 mL) of the filtrate and the resultant filter were retained and frozen at −20 °C until analysis. Total phosphorus in the cell-free medium, and POP, were analysed after defrosting the frozen filtrate and filters and analysed as described above. As the diel light-dark cycle and the daily growth cycle will affect cellular P uptake, all experiments were carried out at the same time of day (08.30), at 15 °C and a light intensity of 50 μmol photons m^−2^ s^−1^. As the experiments were carried out during the day, and for only 4 hours, no cell division occurred.

### Nutrient uptake model

As phytoplankton growth is often limited by nutrient supply, the competitive ability of phytoplankton is affected by their nutrient uptake affinity. Thus, we can think of nutrient uptake affinity as an estimate of their competitive abilities at low nutrient concentrations [39]. Therefore, in order to directly link growth with nutrient uptake, we must first quantify phytoplankton biomass in terms of the amount of limiting nutrient [40]. As phytoplankton have a variable chemical composition in terms of their nutrient content, we assessed nutrient uptake across a range of P concentrations in batch culture as described above. In the nutrient uptake experiments, we evaluated changes to the internal nutrient store of the cells over time. We tested a number of equations including those formulated by [41] and [42]. However, as the uptake parameters used to describe uptake affinity are analogous to those used to describe primary production [43], we found that the expression depicted by [44] and discussed in detail in [40], created the best fit for our data (expression 10). The internal nutrient store is symbolised by Q and was measured in our experiments as μmol P per cell^−1^. Therefore, Q_max_ is the maximum Q and *Q_0_* is the subsistence quota for P, which is the minimum P concentration required for growth. Where Q > Q_0_, there is enough P available for reproduction. When plotting Q over time, Q should increase quasi-linearly at the start. The slope of the initial increase in Q is denoted by μ(∝mol P cell^−1^ min^−1^), which is the initial uptake rate or the change in internal phosphorus concentration over time. The time where P uptake deviates from linearity and becomes independent of time is defined as *T.* This time was derived from expression 11. The curve fitting was obtained using the Levenberg-Marquardt iteration algorithm used for solving generic curve-fitting problems, with Q_0_, ∝ and Q_max_ as variable parameters. Curve fitting was performed in OriginPro 7 (originLabs). This provided a very tight fit to our experimental data.

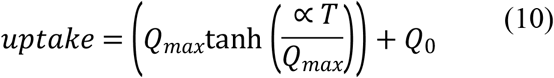

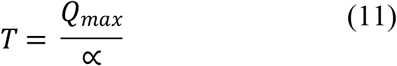

Uptake dynamics were then assessed by plotting initial uptake rates (∝) at the different P concentrations. Uptake was described using the Michalis Menten equation:

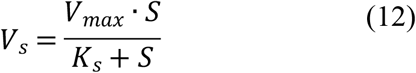

with S being the initial P concentration in the media which declines over time, and V_max_, the maximum uptake rate achieved when S ≥ K_s_. With, K_s_ being the half-saturation constant or the P concentration at which the reaction rate is half of V_max_.

Constants for the models were calculated using the generalised reduced gradient (GRG) non-linear algorithm in Solver in Microsoft Excel.

The maximum growth efficiency was then calculated by:

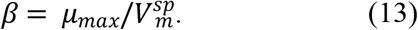

Where 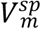, is the maximum specific uptake rate or the ratio of V_max_ to Q_0_ [45].

### Statistical analysis

Statistical procedures were carried out using the Minitab statistical software. Data was initially tested for normality, while some data required log-transformation before statistical analysis. Cell data was analysed using a general linear model (GLM), and nutrient data was analysed using an analysis of variance (ANOVA). P<0.05 was considered significant and variability was measured by standard error of the mean (SEM).

## Results

### Batch Culture

The batch culture experiments were used to calculate the growth rates used in the chemostat experiments. At 2 μM P, growth rates were 0.24 +/− 0.06 for *T. suecia* and 0.22 +/− 0.05 div day^−1^ for *S. Salina,* therefore the flow rate of the chemostat was set at 0.2 div day^−1^. At 0.2 μM P the grown rate was 0.14 +/− 0.04 for *T. suecia* and 0.13 +/− 0.06 div day^−1^ for *S. Salina,* therefore the flowrate of the chemostat was set at 0.1 div day^−1^.

### Chemostat experiments

#### Cell yields

In the 0.2 μM P mono-culture chemostat experiments, both species reached a steady state in terms of cell numbers and biomass after 10 days (Fig. 1a). The green algae *T. suecia* reached a maximum cell density of 2.62 × 10^3^ cell mL^−1^ +/− 1.04 × 10^2^ cells, and the cyanobacterium *S. salina* reached a maximum cell density of 2.09 × 10^5^ +/− 7.03 × 10^3^ cells. When the cell numbers were converted to biomass (μm^3^) (Fig 1.b), total biovolume for *S. salina* was 6.18 × 10^5^ μm^3^ +/− 4.19 × 10^4^, which was significantly higher than that of *T. suecia* 2.12 × 10^5^ μm^3^ +/− 5.85 × 10^3^ (P < 0.01).

**Fig 1.**
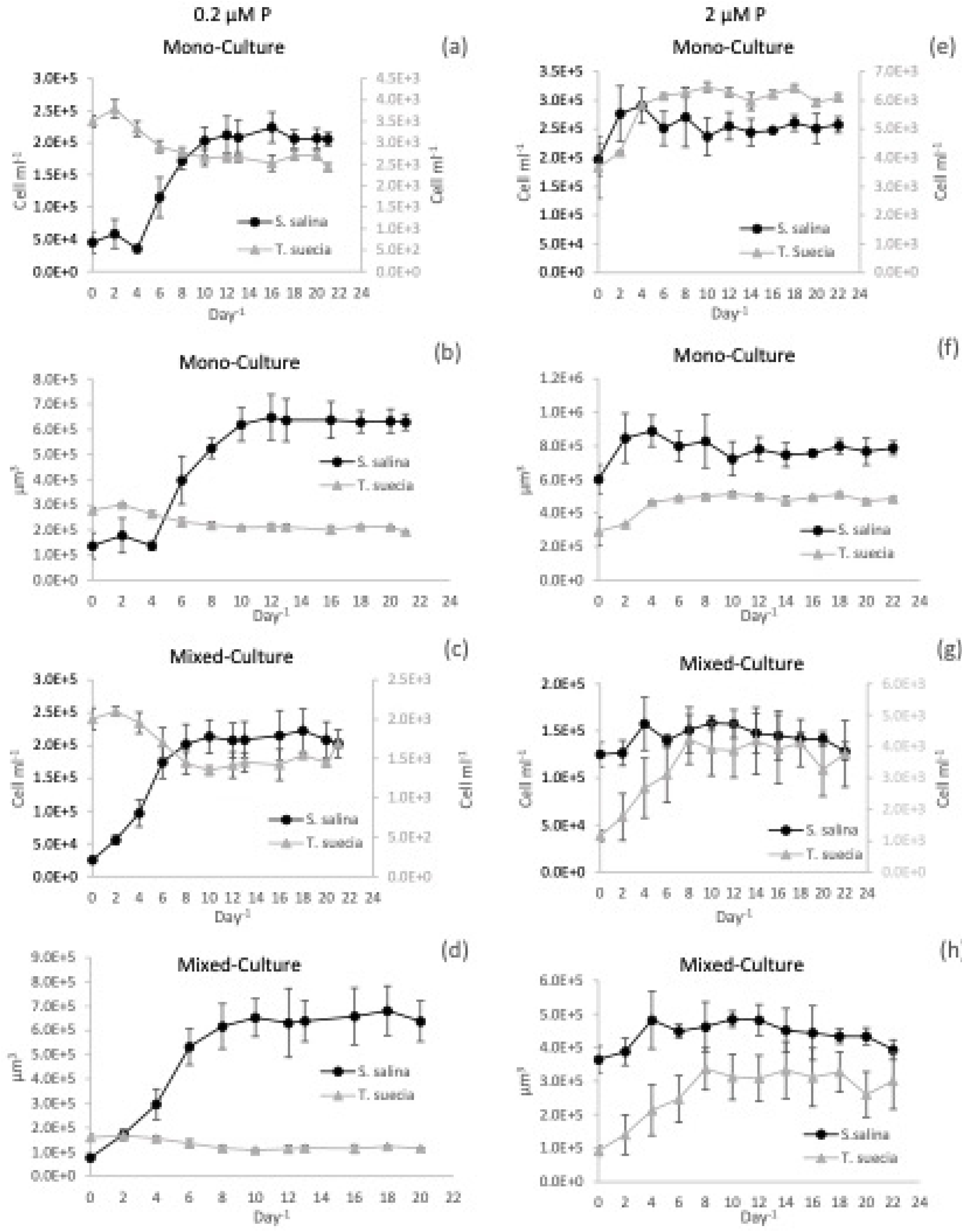
Total biomass for the chemostat experiments over time (Day^−1^) expressed as either cell ml^− 1^ or total biovolume (μm^3^) for the mono-culture and mixed-culture experiments for *S. salina (*•) and *T.suecia* (▲), at 0.2 μM P, Figs (a-d) and 2μM (e-h). Cell numbers are expressed as cell ml^−1^ on the primary axis for *S. salina* and on the secondary axis for *T.suecia* (a, c, e and g). Cell numbers were converted to total biovolume (μm^3^) at 0.2μM P (Figs b and d) and 2μM P (Figs f and h). Error bars represent SEM.

In the mixed chemostat cultures 0.2 μM P, a steady state was reached after 8 days; *T. suecia* reached a maximum cell density of 1.47 × 10^3^ cell mL^−1^ +/− 1.04 × 10^2^ cells and *S. salina* 2.10 × 10^5^ +/− 6.88 × 10^3^ cells (Fig. 1c). When the mixed culture cell numbers were converted to biovolume, there was a large difference between cultures (Fig 1d), where *S. salina* had a biovolume of 6.45 × 10^5^ μm^3^ +/− 2.08 × 10^4^, a value significantly larger than for *T. suecia* at 1.15 × 10^5^ μm^3^ +/− 1.60 × 10^3^ (P < 0.01).

In the chemostat receiving 2 μM P, the mono-culture treatments for both species reached a steady state after 6 days (Fig 1e.). *Tetraselmis suecia* reached a maximum cell density of 6.2 × 10^3^ cell mL^−1^ +/− 1.90 × 10^2^ cells while *S. salina* reached a cell density of 2.54 × 10^5^ +/− 1.02 × 10^4^ cells. When the mixed culture cell numbers were converted to biovolume (μm^3^) (Fig 1f), *S. salina* reached a total biovolume of 8.53 × 10^5^ μm^3^ +/− 2.23 × 10^4^, which was significantly higher than the biovolume of *T. suecia* at 4.93 × 10^5^ μm^3^ +/− 1.51 × 10^4^ (P < 0.01). In the mixed culture 2 μM P chemostat experiments, a steady state in cell numbers was reached after 8 days (Fig 1g), where *T. suecia* reached a maximum cell density of 9.13 × 10^2^ cells mL^−1^ +/− 1.36 × 10^2^ cells and *S. salina* had a cell density of 1.5 × 10^4^ +/− 7.92 × 10^3^ cells mL^−1^. When the mixed cultures cell densities were converted to biovolume, the total biovolume for *S. salina* was 4.48 × 10^5^ μm^3^ +/− 2.96 × 10^4^, which was marginally larger than that of *T. suecia* at 3.11 × 10^5^ μm^3^ +/− 2.42 × 10^4^ (P < 0.05).

**Table 1.** Mean minimum P concentrations (μM P) in the medium for the 0.2 and 2 μM P treatments for *S. salina* and *T suecia.*

|  | 0.2 μM P | 2 μM P |
| --- | --- | --- |
| *S. salina* | 0.027 +/- 0.01 | 0.04 +/- 0.01 |
| *T. suecia* | 0.035 +/- 0.02 | 0.05 +/- 0.02 |

**Table 2.** Initial conditions, uptake parameters values used for expression (10) and output parameters for *T. suecia* and *S. salina.*

| <u>*T. suecia*</u> | 100 $\mu\text{M}$ | 5 $\mu\text{M}$ | 2 $\mu\text{M}$ | 0.2 $\mu\text{M}$ |
| --- | --- | --- | --- | --- |
| $Q_{\max}$ ($\mu\text{mol P}^{-1}$) | $2.71 \times 10^{-6}$ | $2.55 \times 10^{-6}$ | $1.8 \times 10^{-6}$ | $1.04 \times 10^{-6}$ |
| $Q_0$ ($\mu\text{mol P}^{-1}$) | $8.7 \times 10^{-7}$ | $2.75 \times 10^{-7}$ | $4.3 \times 10^{-7}$ | $1.33 \times 10^{-7}$ |
| $\alpha$ ($\mu\text{mol P cell}^{-1} \text{ min}^{-1}$) | $1.8 \times 10^{-8}$ | $1.6 \times 10^{-8}$ | $1.41 \times 10^{-8}$ | $1.40 \times 10^{-8}$ |
| $T$ (min) | 47.45 | 34.2 | 30.38 | 24 |
| <u>*S. salina*</u> |  |  |  |  |
| $Q_{\max}$ ($\mu\text{mol P}^{-1}$) | $6.76 \times 10^{-8}$ | $4.78 \times 10^{-8}$ | $2.43 \times 10^{-8}$ | $1.87 \times 10^{-8}$ |
| $Q_0$ ($\mu\text{mol P}^{-1}$) | $1.19 \times 10^{-8}$ | $1.69 \times 10^{-8}$ | $1.63 \times 10^{-8}$ | $2.85 \times 10^{-9}$ |
| $\alpha$ ($\mu\text{mol P cell}^{-1} \text{ min}^{-1}$) | $5.56 \times 10^{-10}$ | $4.78 \times 10^{-10}$ | $4.58 \times 10^{-10}$ | $4.56 \times 10^{-10}$ |
| $T$ (min) | 71.8 | 35.56 | 26.6 | 18 |

#### Chlorophyll *a*

The co-culture data has not been included in our Chl *a* analysis as we could only perform bulk chlorophyll measurements, so no data is available for the individual species in these experiments. For the 0.2 μM P treatment, Chl *a* concentrations per cell for *T. suecia* remained relatively constant, ranging between 2.02 × 10^−6^ μg cell^−1^ and 4.72 × 10^−6^ μg cell^−1^ with no clear trend (Fig 2a). Cellular Chl *a* concentrations for *S. salina* started high at 9.08 × 10^−7^ μg cell^−1^ before declining to 1.72 × 10^−7^ μg cell^−1^ by day 8, after which concentrations remained steady. When chlorophyll concentrations were calculated relative to biovolume, concentrations were initially significantly higher for *S. salina* (P < 0.01) with a maximum of 2.97 × 10^−4^ μg chl^− 1^μm^−1^ until day 8, after which concentrations became statistically similar between the two species (GLM, P >0.05) (Fig 2c). Chlorophyll *a* concentrations for *T. suecia* fluctuated between a maximum of 5.9 × 10^−5^ μg chl^−1^μm^−1^ and a minimum of 2.54 × 10^−5^ μg chl^−1^μm^−1^.

**Fig 2.**
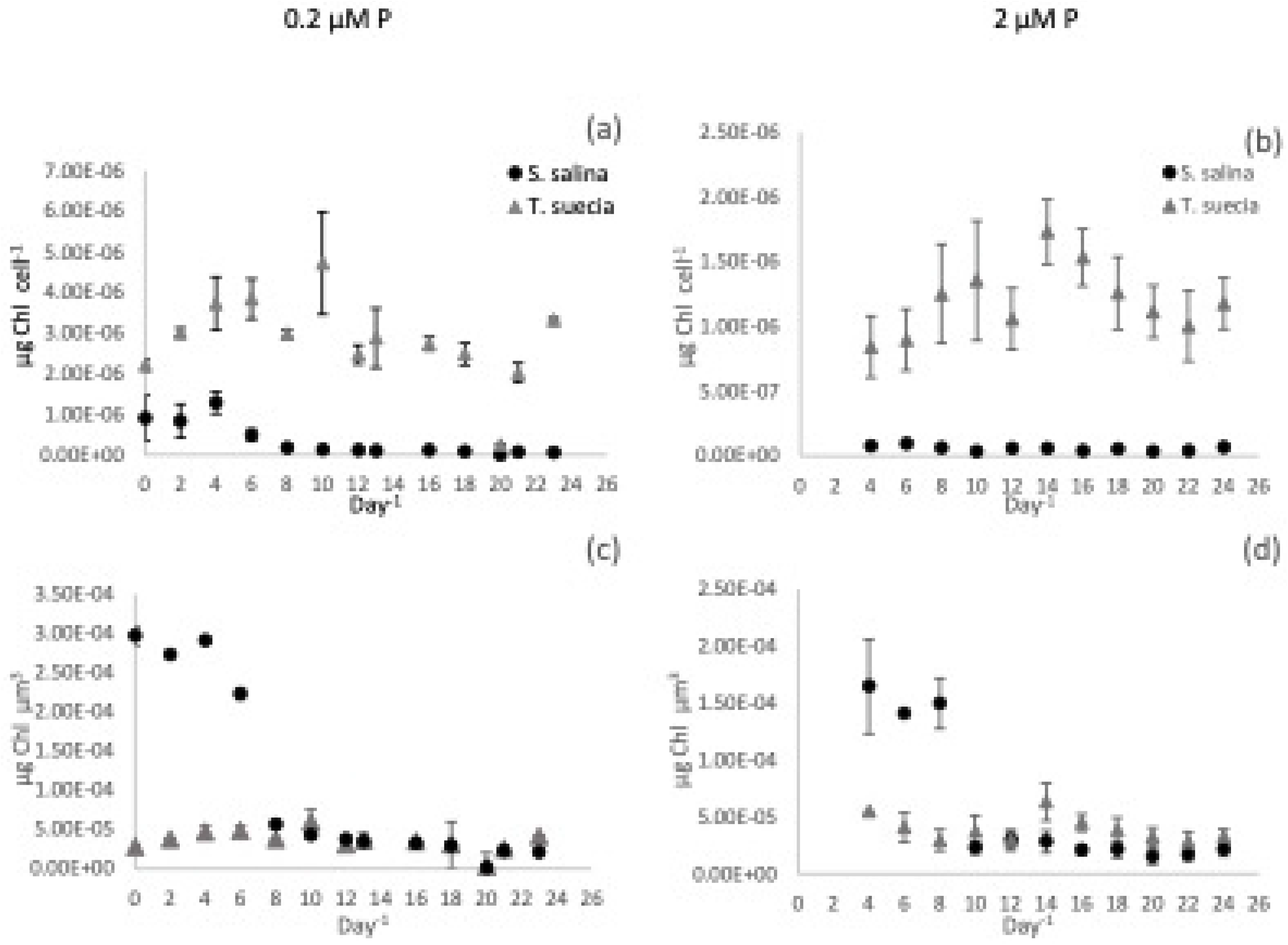
Chlorophyll concentrations per cell^−1^ (a and b) and per unit biovolume (μm^3^) (c and d) per day^−1^ for the mono-culture experiments for *S. salina (*•) and *T.suecia* (▲) at 0.2μM P (a) and 2μM P (b). Error bars represent SEM.

In the 2 μM P treatment, chlorophyll concentrations for *S. salina* ranged from a high of 9.26 × 10^−8^ μg cell^−1^ on day six to a minimum of 3.61 × 10^−8^ μg cell^−1^ on day 10 (Fig 2b). Concentrations for *T. suecia* ranged from a minimum of 8.41 × 10^−7^ μg cell^−1^ on day 4 to a maximum of 1.72 × 10^−6^ μg cell^−1^ on day 14. When adjusted for biovolume (Fig 2d), *S. salina* had a significantly higher Chl *a* concentration (P < 0.01) compared to *T. suecia*, with the exception of days 14-16 when *T. suecia* had a slight but significantly higher Chl *a* content (P < 0.05). The maximum concentration for *S. salina* was 1.65 × 10^−4^ μg chl^−1^μm^−1^ on day 4 and this declined to a minimum of 1.53 × 10^−5^ μg chl^−1^μm^−1^ on day 18. Chlorophyll *a* concentrations for *T. suecia* fluctuated between a maximum of 6.3 × 10^−5^ μg chl^−1^μm^−1^, to a minimum of 2.83 × 10^−5^ μg chl^−1^μm^−1^.

Chlorophyll concentrations are commonly correlated to internal and external phosphorus concentrations [46–49]; however, we found no direct relationship between Chl *a* concentration for either internal cell quota Q or external P concentrations using a Pearson product-moment correlation (P > 0.05).

### Internal P concentrations

For both treatments, the green alga *T. suecia* had a significantly higher internal P pool compared to *S. salina* both when calculated per cell and when adjusted to biovolume (Fig 3). For the 0.2 μM P treatment, concentrations per cell ranged from 2.39 × 10^−6^ μmol P cell^−1^ to 1.41 × 10^−4^ μmol P cell^−1^ for *T. suecia* and 9.07 × 10^−7^ μmol P cell^−1^ to 1.67 × 10^−7^ μmol P cell^−1^ for *S. salina* (Fig 3a). When adjusted for biovolume, and after the cells reached a steady state (day 10), *T. suecia* had a higher internal P concentration compared to *S. salina* at all time points with exception of days 10 and 12, where internal P concentrations were similar (Fig. 3a). The same was observed in the 2 μM P treatment, where *T. suecia* had a significantly higher internal P concentration per biovolume compared to *S. salina* at all time points except at day 0 (P < 0.05). Internal P concentrations ranged from 6.94 ×10^−4^ μM P cell^−1^ to 1.55 × 10^−5^ μM P cell^−1^ for *T. suecia* and 1.2 ×10^−6^ μM P cell^−1^ to 2.84 ×10^−4^ μM P cell^−1^ for *S. salina* (Fig 3d).

**Fig 3.**
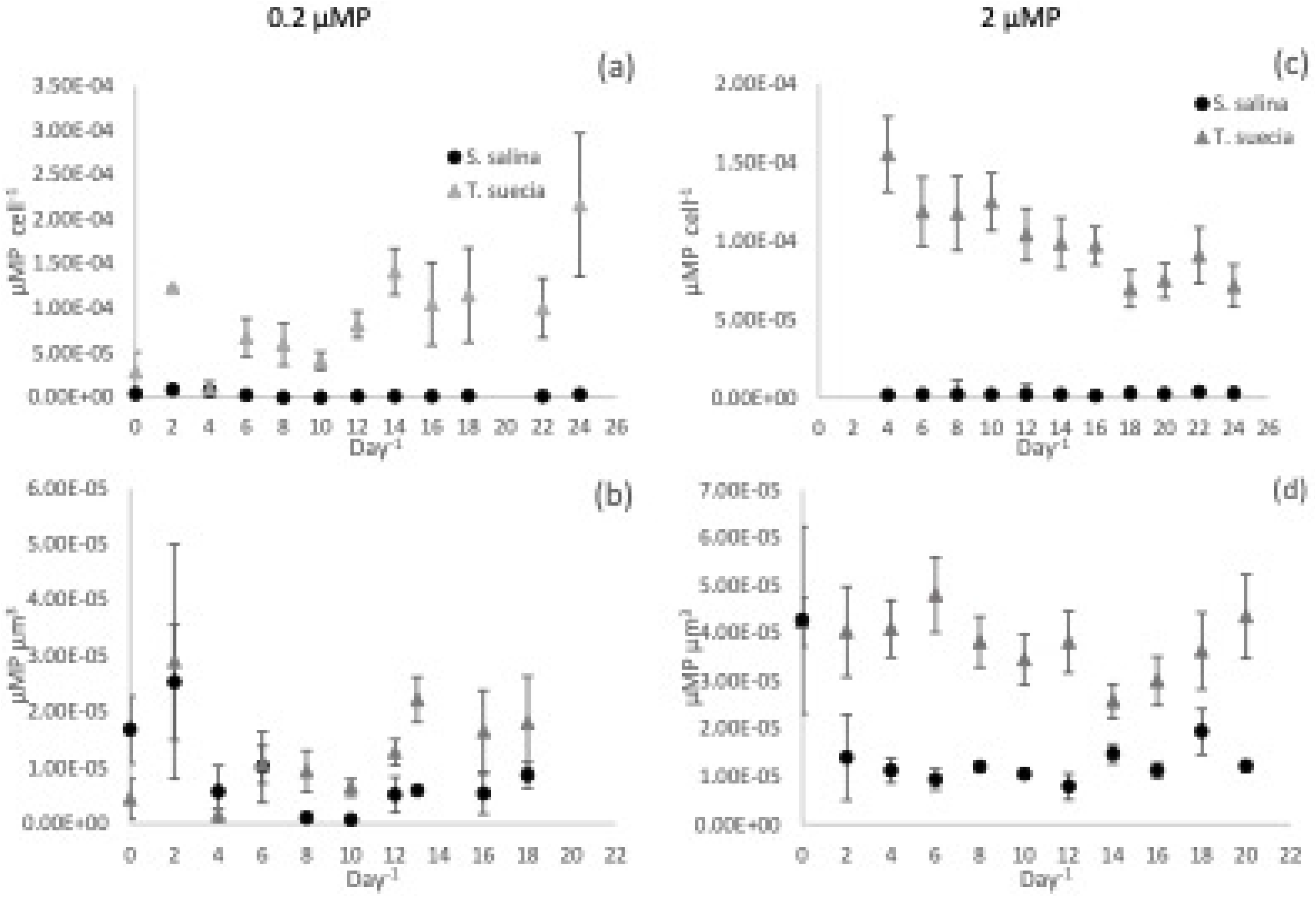
Intracellular phosphorus concentrations per cell^−1^ (a and c) and per unit biovolume (μm^3^) (b and d) per day^−1^ for the mono-culture experiments for *S. salina (*•) and *T.suecia* (▲) at 0.2μM P (a and b) and 2μM P (c and d). Error bars represent SEM.

### External P Concentrations

After initial inoculation, the concentration of dissolved P in the chemostat experiments decreased through uptake for all treatments to low concentrations, although it was never completely depleted from the media. Minimum values are shown in Table 1. These values were not statistically different from one another (P > 0.05).

### Alkaline Phosphatase Activity

Alkaline phosphatase activity (APA) increased over time for both species in all treatments (Fig 4). For the 0.2 μM P treatment, APA activity started at 0.4 for *T. suecia* and 0.6 nmol L^−1^ min^−1^ *S. salina*. The green algae then reached a maximum rate of 5.14 nmol L^−1^ min^−1^, which was significantly higher than for *S. salina* with a rate of 2.9 nmol L^−1^ min^−1^ (P < 0.05). The APA for the co-culture treatment fell between the two individual species at 3.69 nmol L^−1^ min^−1^ (Fig 4a). Enzyme activity was lower in the 2 μM P treatment (Fig 4b), where *T. suecia* reached a maximum of 1.9 nmol L^−1^ min^−1^ while *S. salina* reached a maximum of 1.2 nmol L^−1^ min^−1^ (P < 0.05). The APA in the mixed-culture treatment closely mirrored the trends for *T. suecia*, with a similar maximum at 1.9 nmol L^−1^ min^−1^.

**Fig. 4.**
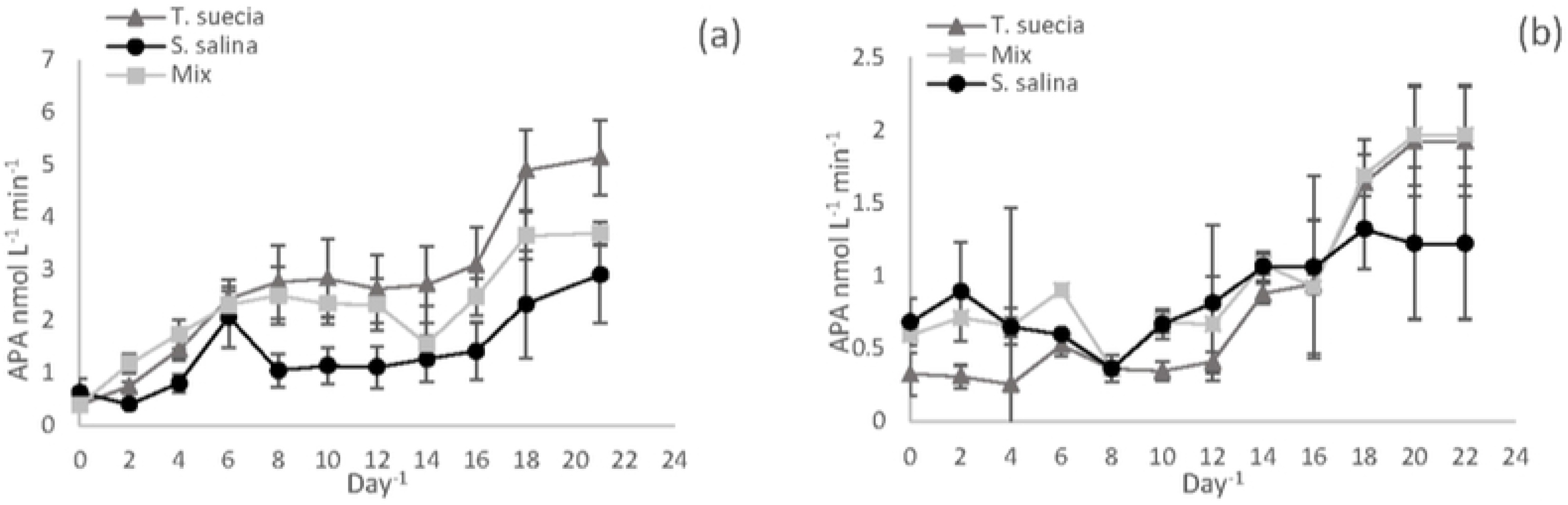
Alkaline Phosphatase activity for the mono-culture experiments for *S. salina(*•) and *T. suecia* (▲) and mixed culture experiments (◻) at 0.2μM P (a) and 2μM P (b). Error bars represent SEM.

### Phosphorus Uptake Following Starvation

The uptake rate of P into the cells was examined at four different dissolved phosphate concentrations following phosphorus starvation. Uptake was measured as μmol P cell^−1^ over the time for the four-hour incubation period. The response seen in the uptake curves (Fig 5) can be described as follows: at time t_0_ the starved cells have an internal phosphorus concentration described by Q_0_ (μmol P^−1^), which is the subsistence quota for P. This is the minimum P concentration required for growth, below this value no growth can occur. Due to this, the uptake curve does not pass through the origin (see Fig 5). As phosphate is re-introduced into the media, the internal P concentration (Q) increases quasi-linearly with a slope denoted by ∝ (μmol P per cell min^−1^). This initial uptake rate is the initial change in internal phosphorus concentration over time. The slope of the curve then decreases progressively until it plateaus. This plateau is described by Q_max_, and at this point the cell is saturated with P. The time at which P uptake deviates from linearity and becomes independent of time is *T* (min).

**Fig 5.**
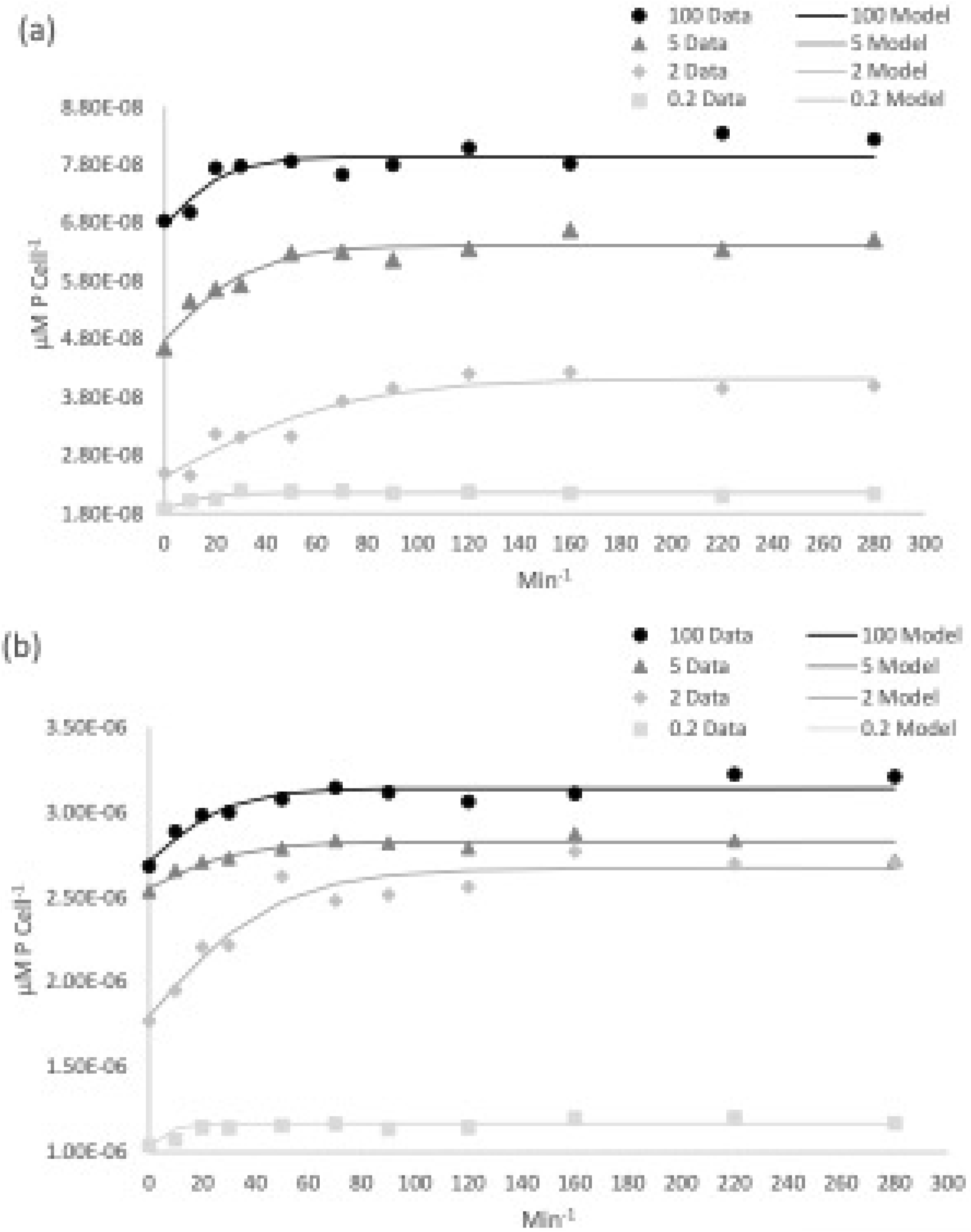
Uptake of intracellular P for *S. salina* (a) and *T. suecia* (b) over a four-hour time course experiment. Experimental data is shown with symbols while the modelled data is shown by lines. Extracellular nutrient concentrations ranged from 100μM P (•), 5 μM P (▲), and 2 μM P (⬪) and 0.2 μM P (◻).

For all phosphate concentrations, most of the uptake occurred within the first hour, after which the uptake began to level off and no further uptake was observed after two hours (Fig 5). Using expression 10, the parameters Q_0_ (subsistence quota), ∝ (initial uptake rate) and Q_max_ (the maximum internal phosphorus concentration) were simultaneously calculated (see Methods) from the experimental data from each phosphate concentration. The parameters derived are shown in Table 2.

The time at which the initial uptake began to slow (*T*) varied between species and phosphate concentrations, and ranged between 71.8 min and 18 min for *S. salina* and 47.45 min and 24 min for *T. suecia*. At the highest phosphate concentrations (100 μM), *T* occurred earlier for *T. suecia* than for *S. salina.* At 5 μM, uptake slowed at a similar time for both species and at 2 and 0.2 μM, uptake slowed first for *S. salina* (Table 2).

We then assessed uptake dynamics by plotting uptake rates (∝) at the different P concentrations. Uptake was described using the Michaelis-Menten equation (expression 12) (Fig 6). *Tetraselmis suecia* had a half saturation constant (K_s_) of 0.4 μM and a maximum uptake rate (V_max_) of 0.16 pmol P cell h^−1^. While *S. salina* had a K_s_ of 0.02 μM and a V_max_ of 0.05 pmol P cell h^−1^. The differences were significant between the two species (P < 0.05). The uptake rate ∝ increased with increasing P concentration until it plateaued at 7 μM for *T. suecia,* after which uptake was at its maximum. For *S. salina* this concentration was 4.3 μM. Michaelis-Menton parameters are summarized in Table 3.

**Table 3.** The maximum uptake (V_max_,) and the half saturation constant (K_s_) described by expression 12.

| | $K_s$ ($\mu\text{M}$) | $V_{\text{max}}$ (pmol P cell $\text{h}^{-1}$) |
| --- | --- | --- |
| *S. salina* | 0.020 | $5 \times 10^{-4}$ |
| *T. suecia* | 0.40 | 0.016 |

**Fig 6.**
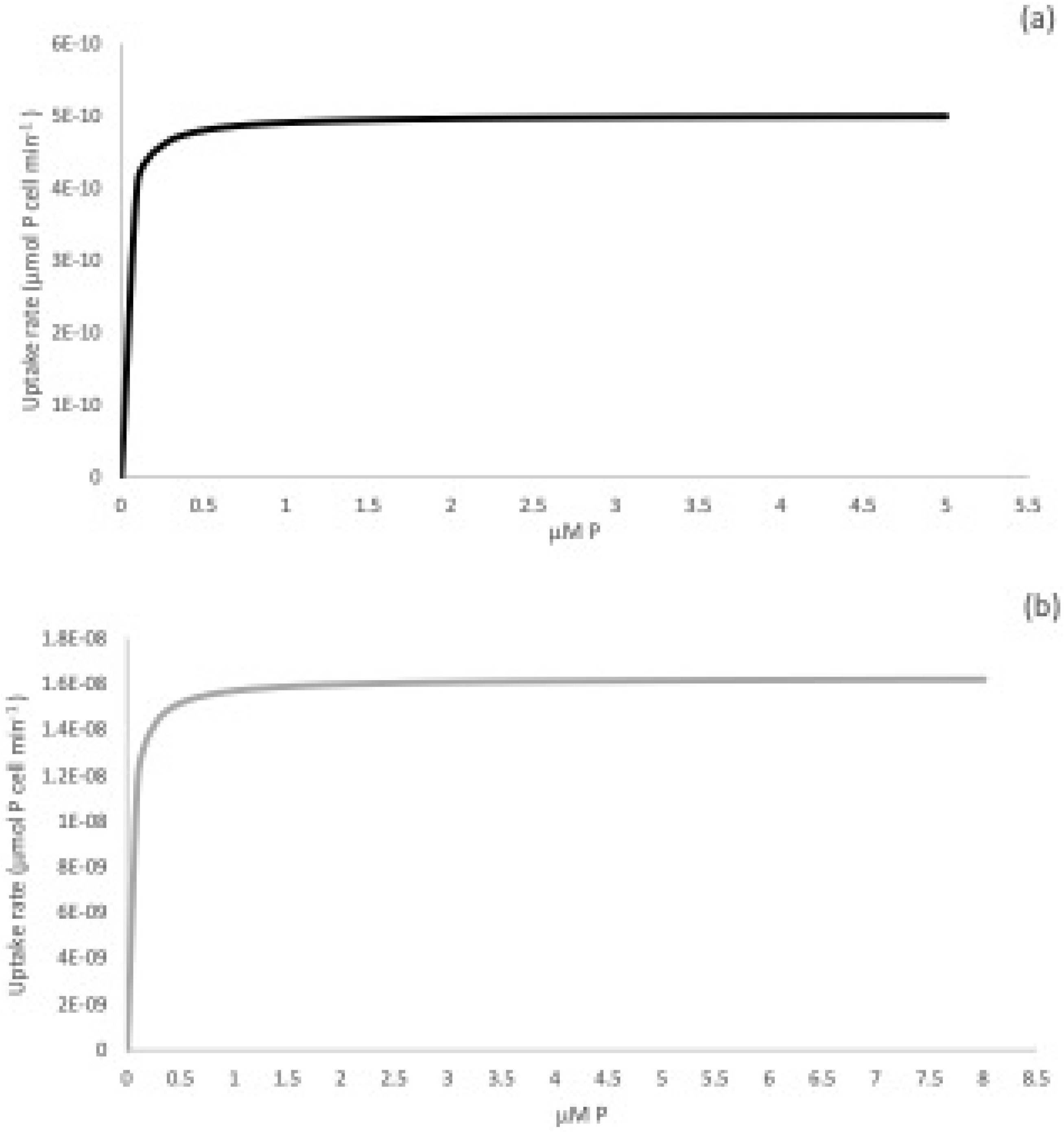
Uptake rates ∝ (μmol P cell min^−1^) plotted against external P concentration (μM) for *S. salina* (a) and *T. suecia* (b). So that fine detail can be observed Figs are only plotted until just after the uptake rate plateaued.

On average, *T. suecia* was 32 times larger in terms of volume than *S. salina*, and owing to its greater cell size, it had a larger internal nutrient store (Q) and initial uptake rate (∝) (Table 2). As we cannot compare parameters between the two species due to their different sizes, we evaluated the efficiencies of nutrient uptake on the basis of their cell subsistence quota (Q_o_). This was done by examining the maximum specific uptake rate 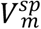, which is the ratio of V_max_ to Q_0._. The 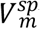 was 8.18 h^−1^ (196.32 d^−1^) for *T. suecia* and 5.68 h^−1^ (136.32 d^−1^) for *S. salina*. These values mean that to maintain their maximum growth rates, *T. suecia* must take up 196.32 +/− 12 times its Q_0_ per day, whilst *S. salina* must take up 136.3 +/−10 times it’s Q_0_ per day. In order to compare the efficiency of P uptake, the maximum growth efficiency *β* was calculated (Equation 13). This value was 1.26 × 10^−1^ +/− 1.2 × 10^−2^ for *T. suecia* and 1.9 × 10^−1^ +/− 1.6 × 10^−2^ for *S. salina.* The higher maximum growth efficiency of the cyanobacteria means that *S. salina* is more efficient at taking up and utilising P and turning it into biomass compared to *T. suecia*.

## Discussion

This study examines the dual hypotheses that phosphorus limitation accounted for the dominance of cyanobacteria before the expansion of algae during mid-Neoproterozoic times, and that this expansion could have resulted from an increase in phosphorus availability. To address these hypotheses, we studied two phosphate-limited conditions meant to simulate both severe phosphorus limitation and phosphorus replete conditions. Chemostats were chosen as they are well suited for the physiological characterisation of microorganisms, especially when investigating the effect of changing cultivation parameters. In this case we explored the role of P concentration on cell performance. The chemostats were fed with fresh medium containing either 0.2 μM P or 2 μM P at dilution rates generating growth rates of 0.1 d^−1^ (for the 0.2 μM P treatment) and 0.2 d^−1^ (for the 2.0 μM P treatment). The concentration of 2 μM P approximates modern deep-water phosphate concentrations [50], while 0.2 μM P has been estimated for deep-water phosphate levels during the Mesoproterozoic Era [21]. In addition, the growth rates in our chemostats can be compared to those measured in the ocean [51,52], where the lower growth rate is typical for algae in modern oligotrophic waters [53]. While the higher growth rate is similar to those found for both algae and cyanobacteria in higher productivity regions, for example those found in the South Atlantic Ocean [54].

—As cells were inoculated into chemostats containing either 0.2 or 2 μM P, nutrients were in excess before the chemostats reached equilibrium. This period of time is reminiscent of an early spring bloom with a plentiful supply of nutrients. After the nutrients were utilised and a steady state was reached, growth rates then reflected the rate of nutrient supply [55] as described by the Monod equation [56,57]. This open system then allows for continuous exponential growth under constant conditions.

### Mono and mixed-culture experiments

We undertook both mono- and mixed-culture experiments in order to examine the full spectrum of physiological behaviour of both species. The mono-culture experiments were used to establish the basic growth parameters for the different species under different levels of phosphate limitation, while the mixed-culture experiments demonstrated how these two organisms compete for phosphate as a limiting nutrient. Theoretically, in the absence of any other interactions or processes such as grazing or cell death, the outcome of competition at a given dilution rate and substrate concentration depends on the relationship between specific growth rate, substrate concentration and internal nutrient store [57], and these specific characteristics of growth vary between species. For this reason, different species will likely should dominate in ecosystems depending on rates of nutrient supply [58–60].

A clear finding from our experiments is that phosphorus loading has the potential to influence the composition of a phytoplankton community. Thus, in our 0.2 μM P mixed culture experiments, the cyanobacterium *S. salina* dominated in biovolume by about a factor of six over the alga *T. suecia* under these, our most phosphate limiting conditions (Fig 1d). Indeed, the maximum cell yield for *S. salina* was unaffected by the mixed community conditions, as it was able to achieve the same peak biomass as it did in mono-culture. However, the total biomass for *T. suecia* in the mixed culture was approximately half when compared to mono-culture. Yet despite having a lower biomass, *T. suecia* was not outcompeted to complete exclusion and it remained a low yet constant background component.

When we increased the nutrient input to the chemostats to 2 μM P, concentrations close to modern deep-water values, the difference in total biomass between species in the mixed culture was reduced compared to the 0.2 μMP treatment (Fig 1h). However, the green alga did not outcompete the cyanobacteria. As neither of the species has been shown to have any allelopathic ability, the greater mean cell and biomass yield that was achieved by *S. salina* at all phosphorus concentrations indicates a more efficient uptake and utilisation of P. This is most likely due to the smaller cell size, giving the cyanobacterium favourable phosphorus acquisition and uptake abilities as discussed in detail below.

### Impact of Cell Size

Many physiological traits such as growth rate, metabolism, light utilisation, access to resources and susceptibility to grazing are significantly correlated to cell size [39]. Therefore, as cell size can affect ecological niches, shape community structure and diversity [24,61,62], it is often termed a ‘master trait’ [43]. In the modern ocean, phytoplankton communities often experience a trade-off based on top-down and bottom-up controls related to cell size, such as nutrient uptake abilities vs size-selected grazing. While nutrient limitation drives communities towards smaller cell sizes, grazing pressure pushes the community towards larger cell sizes [62,63]. This means that phytoplankton communities are not static and will change their composition in response to changing nutrient availability and other environmental factors such as light, temperature and grazing pressure [25]. This will result in communities selecting for a different trait depending on environmental conditions [25,64]. In modern temperate oceans there is a pronounced seasonality in relation to cell size and community structure. In winter, nutrient availability and grazing pressure have little influence over algal community structure as nutrients are plentiful and grazing pressure is low. However, in the spring when nutrient concentrations are still high, the impact of grazing pressure becomes more evident, which results in a population containing larger cells. Our system did not account for grazing pressure, but the higher P input in our high P treatment was better able to support a larger population of the larger *T. suecia* cells. As nutrients become utilised over the summer, nutrient availability becomes more important and the resultant community is comprised of smaller cells. In tropical regions, the stable mixing layer and more constant environmental conditions results in a balance between grazing and nutrient uptake, creating a community with a constant mean cell size throughout the year [63]. Our low P treatment while designed to represent conditions during the Mesoproterozoic and early Neoproterozoic Eras is also reminiscent of conditions experienced in modern oligotrophic waters, where low nutrient availability and the increased importance of nutrient uptake from a limited supply favours a dominance of small organisms such as the picophytoplankton [25,65].

### P Acquisition

When the growth rate and biomass of phytoplankton is restricted by a limiting nutrient, the ability for a species to compete for a limiting resource is an important determinant of the community composition. Different organisms will have different strategies for dealing with P limitation. Such strategies can include different uptake abilities, metabolic restructuring of cellular metabolites [66], use of internal P stores, the utilisation of DOP by hydrolytic enzymes, the substitution of sulphate for phosphate in membrane lipids [67] and the use of alternative low P enzymes [68].

Perhaps the most important mechanism for coping with low P availability is the utilisation of dissolved organic phosphorus (DOP) [18,24]. Whilst phytoplankton have a preference for orthophosphate, they are able to utilise other forms such as DOP by hydrolysing the labile fraction into orthophosphate. The process is facilitated by the enzyme alkaline phosphatase. This enzyme has a wide substrate specificity and hydrolyses ester bonds between P and organic molecules [28], and overall, alkaline phosphatase concentrations have become a proxy for P limitation in phytoplankton communities [69,70]. The production of AP by phytoplankton is regulated by both external and internal P concentrations [71]. In our experiments, the onset of P limitation could be tracked by the increase in APA with declining P concentrations in the media, with APA activity increasing until the cessation of the experiment. Alkaline phosphatase activity was higher in the 0.2 μM P conditions signifying a greater level of nutritional stress. APA also varied between the two species and was higher for *T. suecia* compared to *S. salina,* suggesting *T. suecia* was experiencing a higher level of P stress [72].

The chemical form of phosphorus used in the medium was potassium phosphate, but some chemical forms of organic phosphorus in the aquatic environment, such as phosphonates and phosphites, can only be utilised by bacteria and cyanobacteria [73–75]. As phosphonates account for a significant proportion (25%) of the marine DOP pool, the ability to utilise these alternative sources of phosphorus could provide species such *as S. salina* an advantage during P limitation conditions. As our medium was made using aged natural seawater, we cannot exclude that small concentrations of these organic phosphorus compounds were present in the medium. Also, we do not know if *S. salina,* specifically, can utilise such alternative sources, but closely related marine picocyanobacteria *Synechococcus* and *Prochlorococcus* can use them, and they express *phnD*, the gene encoding the phosphonate binding protein [74,75] while *Prochlorococcus* also possess *ptxD,* the gene encoding phosphite dehydrogenase [76].

In a study by [77], a number of different adaptive strategies were described among freshwater phytoplankton for dealing with variable supplies of phosphorus. In their study, under variable P conditions, species were either described as being velocity-adapted, where high rates of P uptake are employed, or storage-adapted where there is a net accumulation of intra-cellar P. Both species of phytoplankton explored in the current study are capable of luxury P uptake, where uptake and storage of P go beyond the levels of immediate growth [78]. The ability to store phosphorus allows short-term uncoupling of growth rate from both external phosphorus concentrations and uptake. Phosphorus is stored as polyphosphate (polyP), which consists of linear chains of phosphate residues lined by phosphoanhydride bonds [79]. The stored P can contain 1.5-9 times the minimum cell quota (Q_0_) (table 2) and can therefore theoretically sustain 3-4 subsequent doublings without taking up additional P [78,80]. The green alga *T. suecia* had a larger internal P pool and could store more P in relation to its minimum cell quota compared to *S. salina*, both per cell and when adjusted to biovolume (Fig 3b). As the stored P can be used to support population growth for multiple generations after the onset of P limitation [24], this physiological difference could provide *T. suecia* with an important adaption in regions with a variable P supply. When P becomes depleted, it could perhaps allow *T. suecia* to maintain a stable population for longer, making it better adapted to environments with fluctuating P conditions.

### Uptake Kinetics

The uptake rate of DIP by cells is controlled by a number of different constraints. These constraints include the affinity of the enzymes and transporters used to bind the phosphate and deliver it into the cell [24], their density at the cell surface, the ambient P concentration, as well as the cell size and shape of the organism. The organism’s size and shape governs both the cell’s surface to volume ratio and the thickness of the diffusive boundary layer [25]. As P diffuses through an aqueous boundary layer which surrounds the cells before reaching the cell surface [81], larger cells will experience greater diffusion limitation compared to smaller cells [82].

Once P reaches the cell surface it is transported into the cell by binding to uptake proteins. Here, nutrient uptake is described by Michaelis-Menten-like kinetics (equation 12), in which the half-saturation constant, K_s_ provides a measure of the binding affinity of the phosphate uptake system. Possessing a low K_s_ is one of the most important adaptations to low external P concentrations. Transport proteins have K_s_ values that are usually categorised as either low or high affinity [24,29]. The higher affinity proteins have lower values of K_s_, while the lower affinity proteins have higher values. Species which possess a low K_s_ are better adapted physiologically to deal with frequently low or chronically low P concentrations [80].

The maximum potential of a cell to uptake nutrients is described by V_max_, the value of which is related to the number of nutrient uptake sites situated across the membrane. Phytoplankton species that possess high affinity phosphorus uptake systems will, therefore, typically have lower K_*s*_ and V_max_ values that make them much more efficient at low nutrient concentrations [24]. In turn, phytoplankton with low affinity uptake systems have high K_*s*_ and V_max_ values and are better suited to high nutrient concentrations. Variations in V_max_ and K_s_ will act to provide different nutrient uptake strategies [24,29].

The synthesis of high and low-affinity transporters is regulated by the internal cell quota, Q, and the maximum uptake rate, V_max_ [24,83]. The expression of high-affinity transporters occurs when there is a low Q, whereas low-affinity transporters are upregulated when there is a high environmental P concentration, such as after a nutrient pulse, or at such times where a fast response to environmental concentrations would be an advantage [83,84]. A number of marine cyanobacteria have both high and low-affinity transporters [85,86], while eukaryotic equivalents of low-affinity transporters have also been identified, including the P transporter, IPT, and the sodium or sulphate dependent P transporter SPT [24]. Screening of cDNA libraries have revealed only a few eukaryotic high-affinity transporter equivalents [24]. These include the high-affinity transporter (PHO) identified in *Tetraselmis chui* by [87]. This transporter is transcriptionally upregulated under P-limited conditions. Identifying whether *T. suecia* has a similar transporter was beyond the scope of this study, but the low half saturation constant K_s_ suggests that it most likely has a high-affinity transporter similar to that of *T. chui* [87].

An analysis carried out by [24], compiled a list of marine phytoplankton species from the literature with high and low affinity uptake systems and compared physiological parameters. Species ranged from dinoflagellates to cyanobacteria. In this compilation, K_s_ varied from 0.58-2.6 μM and V_max_ from 0.038-6.84 pmol cell h^−1^. Smaller K_s_ values are typical for other picocyanobacteria, such as those described by [88] for *Synechococcus* and *Prochlorococcus*, in the North Atlantic, where K_s_ values ranged from 8 × 10^−4^ − 0.032 μM [88]. Such differences in phosphorus kinetics are commonly considered to be caused by size-specific differences, e.g. large surface area to volume ratios and smaller diffusive boundary layers, which is why smaller cells are often selected in nutrient-limited environments [25]. The low K_s_ values of 0.02 and 0.40 μM calculated for *S. salina* and *T. suecia* respectively, indicate that both have a high affinity for P. However, the lower V_max_ of 5×10^−4^ pmol cell h^−1^ calculated for *salina* suggests a greater affinity for P compared to *T. suecia*. As the affinity of phytoplankton for nutrients is related to their competitive abilities, owing to its smaller cell size and resultant greater affinity for P we would expect *S. salina* to consistently outcompete *S suecia* in P depleted environments.

As the green algae *T. suecia* was on average 32 times larger in biovolume than the cyanobacteria *S. salina*, we calculated the maximum specific uptake rate, 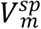 and the the maximum growth efficiency (*β*). These parameters provide an estimate of how proficiently each cell takes up and utilises P regardless of cell size. The 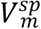 can be used to evaluate the nutrient utilisation efficiency by examining many times per day a cell must fill its subsistence quota, Q_0_, in order to maintain it maximum specific growth rate. While, (*β*) is used to compare the efficiency of P uptake in proportion to growth. *Tetraselmis suecia* had a greater maximum specific uptake of 8.18 h^−1^ (196 d^−1^) compared to 5.68 h^−1^ (136 d^−1^) for *S. salina.* So, in order to maintain their maximum growth rates *T. suecia* must take up 196 times its Q_0_ per day, whilst *S. salina* must take up 136 times it’s Q_0_ per day. This indicates that *T. suecia* has a higher P demand compared to *S. salina,* and therefore it would be unlikely to dominate when P is deficient. Our 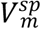value for *T. suecia* (196 d^−1^) is lower that the value calculated for the closely related species *Tetraselmis subcordiformis* by [45], which was 20 h^−1^ (480 d^−1^), but this could be explained by the slightly larger size of *T. subcordiformis* compared to *T. suecia*. However, our 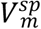 value for *T. suecia* is similar to those calculated for the coccolithophore *Emiliania huxleyi* at 8.90 h^−1^ (214 d^−1^) and the diatom *Thalassiosira pseudonana* 7.80 h^−1^ (187 d^−1^) [89]. Having a low 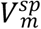 appears to be an important adaptation to P limited conditions. We would therefore, expect species that live in oligotrophic conditions to have a lower 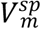 and thus adapted to have a higher capacity for nutrient utilisation, compared to those living in nutrient replete environments.

*Synechocystis salina* had a greater maximum growth efficiency *β*, indicating that it has a lower requirement for P and can reach its maximum specific growth rate at a lower phosphorus concentration [80]. However, growth efficiency is likely to be affected by environmental variables such as temperature, the diel light-dark cycle, irradiance conditions and the associated daily growth cycle [45]. It is common that P uptake rates increase during the day due to the higher demand for P for photosynthetic biomass production. Kinetic analysis of uptake parameters by other authors [24,91,92] have indicated that diel changes in uptake rates will alter the V_max_ but will have no impact on the K_s_ value, suggesting that nutrient uptake rates can fluctuate, but the overall affinity for nutrients remains fixed. Environmental induced fluctuations in the maximum nutrient uptake rate by the more efficient or superior competitor will lead to fluctuations in the ambient nutrient concentration which could positively or negatively impact competing species and thus community compositions. So, if the maximum nutrient uptake rate for *S. salina* declined due to unfavourable environmental conditions, such as altered light and/or temperature regimes, it would make more P available for *T. suecia* or another competing organisms, allowing for an increase in biomass by these competing organisms.

### Impact of P Limitation

Phosphorus deficiency limits phytoplankton productivity by disrupting electron transport to photosystem I (PSI) [94], thus reducing intracellular concentrations of compounds such as ATP, NADPH, nucleic acids, sugar phosphates and phospholipids, all of which are essential in chlorophyll production and ultimately photosynthesis [95]. Chlorophyll *a* is the major photosynthetic pigment of most phytoplankton species and can be used as an index for primary production rate and standing biomass abundance in aquatic ecosystems [93]. The relationship between Chl *a* and phosphorus is a fundamental relationship, with Chl *a* concentrations often having a positive log linear function of total dissolved organic P in both coastal marine [46,47] and fresh water environments [48,49]. However, work by [96] indicated that the chlorophyll *a* content of phytoplankton is not directly related to the external nutrient concentration but rather the internal cell quota, Q. In our study, P concentrations had no impact on Chl *a* concentration which remained constantly low. However, we could also not find a relationship between internal cell quota, Q and Chl *a* concentration for either species. As Chl *a* concentrations are regulated by the balance of energy supplied to PSII and by light harvesting, plus the energy demand for photosynthesis and growth, we can hypothesise that despite P starvation, photosynthesis rates and pigment synthesis remained stable and cell division rates were low enough to maintain a stable chlorophyll concentration [97,98]. This was observed by [99,100] for *Dunaliella tertiolecta* which had a growth rate of 0.24 div day^−1^ in P replete conditions, which is lower than the growth rates used in this study.

### Geobiological implications and concluding remarks

We conducted a series of chemostat-based growth experiments to test the competition for phosphorus between the cyanobacterium *S. salina* and the eukaryotic alga *T. suecia*, representing both different *Domains* in the tree of life and, importantly, different cell sizes as would be typical when comparing cyanobacteria to eukaryotic algae. Ultimately, our experiments were designed to test whether phosphorus availability and changing phosphorus concentrations could explain the history of cyanobacterial versus algal dominance through the Proterozoic Eon.

The organisms we studied are found to coexist in nature, and our chemostat experiments were conducted at growth rates that may be considered typical for these types of organisms in nature. Our experiments explored how these two organisms responded both individually and in co-culture to a range of nutrient limitations. In one case, the organisms were fed with phosphate at near-modern bottom water concentrations, and in another case, they were fed with much more limited phosphorus concentrations, believed to represent bottom water levels from the Mesoproterozoic and early Neoproterozoic Rras. In addition, we conducted a series of batch experiments at different phosphorus levels, and under cell starvation, to calculate the growth kinetics of these organisms relative to phosphorus concentration, as well as how each of these organisms internally stores phosphorus. In addition, APA was monitored in our chemostat experiments as an independent measure of phosphorus stress.

Our results showed that that the competitive outcome of cyanobacteria and eukaryotic algae are heavily influenced by phosphorus concentrations. The cyanobacterium outcompeted the alga at both low and high P treatments, yet the eukaryotic algae were never completely excluded even in the low P treatment. In the higher P treatment, *T. suecia* was able to increase its biomass but was still unable to outcompete the cyanobacteria. This suggests that no matter the P concentration, *S. salina* was consistently the superior competitor for P. This is supported by the low half saturation constant (K_s)_ and uptake rate (V_max_) calculated for *S. salina,* indicating that it possess a higher affinity for P compared to *T. suecia*. This, combined with its low maximum specific uptake rate 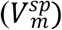 and higher maximum growth efficiency (*β*), further demonstrates that the cyanobacterium has a higher capacity for nutrient utilisation. Despite also having a high affinity for P, the alga *T, suecia* was physically inhibited by its larger cell size and thus, experienced greater nutritional stress, highlighted by its enhanced alkaline phosphatase activity.

Ultimately, our results reinforce already existing ideas that nutrient availability can have an important bearing on the dominant cell size of phototrophs in nature [25,82,101–104]. Clearly, the cyanobacteria *S. salina* could outcompete the eukaryotic alga *T. suecia* under severe nutrient limitation and all of the kinetic parameters we determined (Ks, V_max_, 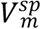 and *β*), support this observation. If *S. salina* represents a typical cyanobacteria of its size, and *T. suecia* represents a typical alga of its size, then our observations are consistent with the idea that phosphorus limitation before the “rise of algae” could have favoured a cyanobacteria-dominated ecosystem. Still, even under severe nutrient limitation, the eukaryotic alga we explored was not excluded from our co-culture chemostats. This would be consistent with observations from the modern ocean where small phototroph sizes are selected under nutrient limitation [25,62,82], but there are no known areas in the modern ocean, even under the most extreme nutrient limitation, where eukaryotic algae are excluded [105–107]. Therefore, our results do not completely match with the biomarker record before about 750 Ma, where steranes are not found, suggesting that eukaryotic algae were of minimal importance in the global marine ecosystem. Indeed, steranes are absent even under a variety of depositional conditions (represented by sedimentary rocks of varying organic matter content) likely representing different degrees of nutrient availability [2,3,108–113].

Our results, would, however be consistent with the finding of various acritarchs through the late-Neoproterozoic and the Mesoproterozoic Eras that are believed to be resting stages of eukaryotic algae [114–118]. Thus, our results would be consistent with cyanobacterial dominance, but not eukaryotic algal exclusion in the Proterozoic oceans. Our results, however, are also consistent with the idea that the “rise of algae” could have been initiated by enhanced phosphorus availability. Indeed, there are some suggestions that phosphorus may have become more available through the Neoproterozoic Era as discussed above. We also note that other aspects of the marine ecosystem were evolving through the same time. For example, grazing ciliates began to occupy the oceans [2,119], and these could have selectively grazed the small-sized cyanobacteria [120–122], providing another avenue towards eukaryotic algal dominance in the Neoproterozoic Era.

Therefore, our results are consistent with enhanced phosphorus availability leading to the “rise of algae” through the Neoproterozoic Era, but our results do not exclude other viable hypotheses for algal dominance in the marine ecosystem. Our results also do not cover the possible role of picoeukaryotes [106,123] (< 2-3 μm in diameter) and their potential ability to have competed with cyanobacteria in the Mesoproterozoic oceans. Indeed, in modern marine environments picoeukaryotic phototrophs prove adept at competing with cyanobacteria under nutrient limitation [103,106,123], although we are unaware of any experiments where phototrophic pico and nano plankton have been compared in their ability to compete with nutrients in the manner that we have presented here.

## Acknowledgements

We wish to acknowledge Heidi Grøn Jensen and Erik Lausen for expert help in the lab.

## References

1. Sánchez-Baracaldo P, Raven JA, Pisani D, Knoll AH. Early photosynthetic eukaryotes inhabited low-salinity habitats. Proc Natl Acad Sci U S A. 2017;114: E7737–E7745. doi:10.1073/pnas.1620089114

2. Brocks JJ, Jarrett AJM, Sirantoine E, Hallmann C, Hoshino Y, Liyanage T. The rise of algae in Cryogenian oceans and the emergence of animals. Nature. 2017; 548: 578–581. doi:10.1038/nature23457

3. Zumberge JA, Rocher D, Love GD. Free and kerogen-bound biomarkers from late Tonian sedimentary rocks record abundant eukaryotes in mid-Neoproterozoic marine communities. Geobiology. 2019; 18(3): 326–347. doi:10.1111/gbi.12378

4. Reinhard CT, Planavsky NJ, Robbins LJ, Partin CA, Gill BC, Lalonde S V., et al. Proterozoic ocean redox and biogeochemical stasis. Proc Natl Acad Sci U S A. 2013;110: 5357–5362. doi:10.1073/pnas.1208622110

5. Shields-Zhou G, Och L. The case for a Neoproterozoic Oxygenation Event: Geochemical evidence and biological consequences. GSA TODAY. 2011;21. doi:10.1130/GSATG102A.1

6. Gernon TM, Hincks TK, Tyrrell T, Rohling EJ, Palmer MR. Snowball Earth ocean chemistry driven by extensive ridge volcanism during Rodinia breakup. Nat Geosci. 2016; 9: 242–248. doi:10.1038/ngeo2632

7. Planavsky NJ, Rouxel OJ, Bekker A, Lalonde S V., Konhauser KO, Reinhard CT, et al. The evolution of the marine phosphate reservoir. Nature. 2010; 467: 1088–1090. doi:10.1038/nature09485

8. Reinhard CT, Planavsky NJ, Gill BC, Ozaki K, Robbins LJ, Lyons TW, et al. Evolution of the global phosphorus cycle. Nature. 2017;541: 386–389. doi:10.1038/nature20772

9. Brocks JJ. The transition from a cyanobacterial to algal world and the emergence of animals. Emerg Top Life Sci. 2018; 2 (2): 181–190. doi:10.1042/ETLS20180039

10. Canfield DE, Poulton SW, Narbonne GM. Late-Neoproterozoic deep-ocean oxygenation and the rise of animal life. Science. 2007; 315: 92–95. doi:10.1126/science.1135013

11. Canfield DE, Teske A. Late Proterozoic rise in atmospheric oxygen concentration inferred from phylogenetic and sulphur-isotope studies. Nature. 1996; 382: 127–132. doi:10.1038/382127a0

12. Scott C, Lyons TW, Bekker A, Shen Y, Poulton SW, Chu X, et al. Tracing the stepwise oxygenation of the Proterozoic ocean. Nature. 2008; 452: 456–459. doi:10.1038/nature06811

13. Och LM, Shields-Zhou GA. The Neoproterozoic oxygenation event: Environmental perturbations and biogeochemical cycling. Earth-Science Reviews. Elsevier; 2012. pp. 26–57. doi:10.1016/j.earscirev.2011.09.004

14. Pogge Von Strandmann PAE, Stüeken EE, Elliott T, Poulton SW, Dehler CM, Canfield DE, et al. Selenium isotope evidence for progressive oxidation of the Neoproterozoic biosphere. Nat Commun. 2015; 6: 1–10. doi:10.1038/ncomms10157

15. Allen JF, Thake B, Martin WF. Nitrogenase Inhibition Limited Oxygenation of Earth’s Proterozoic Atmosphere. Trends Plant Sci. 2019; 24: 1022–1031. doi:10.1016/j.tplants.2019.07.007

16. Ratti S, Knoll AH, Giordano M. Grazers and Phytoplankton Growth in the Oceans: An Experimental and Evolutionary Perspective. PLoS One. 2013; 8: 1–16. doi:10.1371/journal.pone.0077349

17. Porter S. The rise of predators. Geology. 2011; 36(6): 607–608. doi:10.1130/focus062011.1

18. Karl DM. Microbially Mediated Transformations of Phosphorus in the Sea: New Views of an Old Cycle. Ann Rev Mar Sci. 2014; 6: 279–337. doi:10.1146/annurev-marine-010213-135046

19. Falkowski PG, Raven JA. Aquatic photosynthesis. 2nd Edition. In: Princeton University Press [Internet]. 2007 [cited 10 Dec 2019] p. 484. Available: https://press.princeton.edu/titles/8337.html

20. Holland HD. The chemical evolution of the atmosphere and oceans. Princeton University Press; 1984. ISBN: 9780691023816

21. Reinhard CT, Planavsky NJ, Gill BC, Ozaki K, Robbins LJ, Lyons TW, et al. Evolution of the global phosphorus cycle. Nature. 2017; 541: 386–389. doi:10.1038/nature20772

22. Laakso TA, Schrag DP. Regulation of atmospheric oxygen during the Proterozoic. Earth Planet Sci Lett. 2014; 388: 81–91. doi:10.1016/j.epsl.2013.11.049

23. Derry LA. Causes and consequences of mid-Proterozoic anoxia. Geophys Res Lett. 2015; 42: 8538–8546. doi:10.1002/2015GL065333

24. Lin S, Litaker RW, Sunda WG. Phosphorus physiological ecology and molecular mechanisms in marine phytoplankton. Journal of Phycology. Blackwell Publishing Inc.; 2016. pp. 10–36. doi:10.1111/jpy.12365

25. Chisholm SW. Phytoplankton Size. Primary Productivity and Biogeochemical Cycles in the Sea. Springer US; 1992. pp. 213–237. doi:10.1007/978-1-4899-0762-2_12

26. Bjerrum CJ, Canfield DE. Ocean productivity before about 1.9 Gyr ago limited by phosphorus adsorption onto iron oxides. Lett to Nat. 2002; 417: 159–162. Available: www.nature.com

27. Escaravage V, Prins TC, Smaal AC, Peeters JCH. The response of phytoplankton communities to phosphorus input reduction in mesocosm experiments. J Exp Mar Bio Ecol. 1996; 198: 55–79. doi:10.1016/0022-0981(95)00165-4

28. Cembella AD, Antia NJ, Harrison PJ, Rhee GY. The utilization of inorganic and organic phosphorous compounds as nutrients by eukaryotic microalgae: A multidisciplinary perspective: Part 2. Crit Rev Microbiol. 1984; 11: 13–81. doi:10.3109/10408418409105902

29. Cáceres C, Spatharis S, Kaiserli E, Smeti E, Flowers H, Bonachela JA. Temporal phosphate gradients reveal diverse acclimation responses in phytoplankton phosphate uptake. ISME J. 2019; 13: 2834–2845. doi:10.1038/s41396-019-0473-1

30. Blossom HE, Rasmussen SA, Andersen NG, Larsen TO, Nielsen KF, Hansen PJ. Prymnesium parvum revisited: Relationship between allelopathy, ichthyotoxicity, and chemical profiles in 5 strains. Aquat Toxicol. 2014; 157: 159–166. doi:10.1016/J.AQUATOX.2014.10.006

31. LeGresley M, McDermott G. Counting chamber methods for quantitative phytoplankton analysis—haemocytometer, Palmer-Maloney cell and Sedgewick-Rafter cell. 55th ed. In: Karlson B, Cusack C, Bresnan E, editors. Microscopic and molecular methods for quantitative phytoplankton analysis. 55th ed. UNESCO (IOC Manuals and Guides); 2010. pp. 25–30.

32. Shishlyannikov SM, Zakharova YR, Volokitina NA, Mikhailov IS, Petrova DP, Likhoshway Y V. A procedure for establishing an axenic culture of the diatom Synedra acus subsp. radians (Kütz.) Skabibitsch. from Lake Baikal. Limnol Oceanogr Methods. 2011; 9: 478–484. doi:10.4319/lom.2011.9.478

33. Hillebrand H, Sommer U. The nutrient stoichiometry of benthic microalgal growth : Redfield proportions are optimal. Limnol Oceanogr. 1999; 44: 440–446.

34. Fehling J, Davidson K, Bates SS. Growth dynamics of non-toxic Pseudo-nitzchia delicatissima and toxic P.seriata Bacillariophyceae) under simulated spring and summer photoperiods.+ Harmful Algae. 2005; 4: 763–769.

35. Flynn KJ, Davidson K, Leftley JW. Carbon-nitrogen relations during batch growth of Nannochloropsis oculata (Eustigmatophyceae) under alternating light and dark. J Appl Phycol. 1993; 5: 465–475. doi:10.1007/BF02182739

36. Lonborg C, Davidson K, Alvarez-Sagado XA, Miller AEJ. Bioavailability and bacterial degradation rates of dissolved organic matter in a temperate coastal area during an annual cycle.+ Mar Chem. 2009; 113: 219–226.

37. Hansen HP, Koroleff F. Determination of Nutrients. 3rd ed. In: Grasshof K, Kermling K, Ehrhardt M, editors. Methods of seawater analysis. 3rd ed. Germany: Wiley-VCH; 1999. p 159–228.

38. Ohman MD. Predation on planktonic protists assessed by immunochemical assays. In Kemp PF, Sherr BF, Sherr EB, Cole JJ., editors. Handbook of methods in Aquatic Microbial Ecology. Taylor & Francis; 1994. p 1496–1497.

39. Lindemann C, Fiksen Ø, Andersen KH, Aksnes DL. Scaling Laws in Phytoplankton Nutrient Uptake Affinity. Front Mar Sci. 2016; 3: 26. doi:10.3389/fmars.2016.00026

40. Tett P, Droop MR. Cell quota models and plantonic primary production. In: Wimpenny JW, editor. CRC Handbook of Laboratory Model Systems for Microbial Ecosystems, Volume II. CRC Press; 1988. p 177–234.

41. Monod J. Recherches sur la croissance des cultures bacteriennes. Hermann and Cie. 1942. p 911

42. Dugdale RC. The Sea Marine Modelling. In: Goldberg, Edward D., McCave, I. N., O’Brien, J. J., Steele JH, editors. 6th ed. New York: John Wiley & Sons; 1977.

43. Litchman E, Klausmeier CA. Trait-Based Community Ecology of Phytoplankton. Annual Review of Ecology, Evolution, and Systematics. 2008: 39: 615–639. doi:10.1146/annurev.ecolsys.39.110707.173549

44. Jassby AD, Platt T. Mathematical formulation of the relationship between photosynthesis and light for phytoplankton. Limnol Oceanogr. 1976; 21: 540–547. doi:10.4319/lo.1976.21.4.0540

45. Chunrong N, Shuanglin D. Comparative Studies on Phosphorus Uptake and Growth Kinetics of the Microalga *Tetraselmis subcordiformis* and the Macroalga *Ulva pertusa*. J Ocean Univ. China 2004; 3: 56–59. https://doi.org/10.1007/s11802-004-0009-8

46. Giovanardi F, Tromellini. E. Statistical Assessment of Trophic Conditions. Application of the OECD Methodology to the Marine Environment. Proc. Int. Conf. Marine Coastal Eutrophication. Sci Total Env. 1992; 211–233.

47. Vollenweider RA, Rinaldi A, Montanari G. Eutrophication, structure and dynamics of a marine coastal system: results of ten-year monitoring along Emilia-Romagna coast (Northwest Adriatic Sea). In: Vollenweider RA, R. Marchetti, Viviani R, editors. Marine Coastal Eutrophication: Proceedings of an International Conference. Elsevier; 1990. p. 63–106.

48. Filstrup CT, Downing JA. Relationship of chlorophyll to phosphorus and nitrogen in nutrient-rich lakes. Inl Waters. 2017;7: 385–400. doi:10.1080/20442041.2017.1375176

49. Håkanson L, Eklund JM. Relationships Between Chlorophyll, Salinity, Phosphorus, and Nitrogen in Lakes and Marine Areas. J Coast Res. 2010; 26: 412–423. doi:10.2112/08-1121.1

50. Bender ML. Tracers in the Sea. BioScience, 1984; 34,(7), p 452, https://doi.org/10.2307/1309641

51. Goldman JC, McCarthy JJ, Peavey DG. Growth rate influence on the chemical composition of phytoplankton in oceanic waters. Nature. 1979; 279: 210–215. doi:10.1038/279210a0

52. Eppley RW. Relations between nutrient assimilation and growth in phytoplankton with a brief review of estimates of growth rate in the ocean. Can J Fish. 1981. 201: p251–263

53. Jackson GA. Phytoplankton growth and Zooplankton grazing in oligotrophic oceans. Nature. 1980; 284: 439–441. doi:10.1038/284439a0

54. Zubkov M V. Faster growth of the major prokaryotic versus eukaryotic CO_2_ fixers in the oligotrophic ocean. Nat Commun. 2014; 5. doi:10.1038/ncomms4776

55. Droop MR. Vitamin B12 and Marine Ecology. IV. The Kinetics of Uptake, Growth and Inhibition in *Monochrysis Lutheri*. J Mar Biol Assoc United Kingdom. 1968; 48: 689–733. doi:10.1017/S0025315400019238

56. Monod J. The growth of bacterial cultures. A. Rev. Microbiol. 1949; 3: 371–394.

57. Wides A, Milo R. Understanding the Dynamics and Optimizing the Performance of Chemostat Selection Experiments. 2018. arXiv. 2018; (arXiv:1806.00272) https://arxiv.org/abs/1806.00272

58. MacIntyre S. Turbulent mixing and resource supply to phytoplankton. In: Jörg Imberger, editor. Physical Processes in Lakes and Oceans. American Geophysical Union (AGU); 1998. pp. 561–590. doi:10.1029/ce054p0561

59. Paczkowska J, Rowe OF, Figueroa D, Andersson A. Drivers of phytoplankton production and community structure in nutrient-poor estuaries receiving terrestrial organic inflow. Mar Environ Res. 2019; 151: 104778. doi:10.1016/j.marenvres.2019.104778

60. Agawin NSR, Duarte CM, Agustí S. Nutrient and temperature control of the contribution of picoplankton to phytoplankton biomass and production. Limnol Oceanogr. 2000; 45: 591–600. doi:10.4319/lo.2000.45.3.0591

61. Ploug H, Stolte W, Epping EHG, Jørgensen BB. Diffusive boundary layers, photosynthesis, and respiration of the colony-forming plankton algae, *Phaeocystis* sp. Limnol Oceanogr. 1999; 44: 1949–1958. doi:10.4319/lo.1999.44.8.1949

62. Irwin AJ, Finkel Z V, Schofield OME, Falkowski PG. Scaling-up from nutrient physiology to the size-structure of phytoplankton communities. J Plank Res. 2006; 28(5): 159–471. doi:10.1093/plankt/fbi148

63. Acevedo-Trejos E, Brandt G, Bruggeman J, Merico A. Mechanisms shaping size structure and functional diversity of phytoplankton communities in the ocean. Sci Rep. 2015; 5: 1–8. doi:10.1038/srep08918

64. Mena C, Reglero P, Hidalgo M, Sintes E, Santiago R, Martín M, et al. Phytoplankton community structure is driven by stratification in the oligotrophic Mediterranean Sea. Front Microbiol. 2019; 10: 1698. doi:10.3389/fmicb.2019.01698

65. Arin L, Anxelu X, Estrada M. Phytoplankton size distribution and growth rates in the Alboran Sea (SW Mediterranean): short term variability related to mesoscale hydrodynamics. J Plankton Res. 2002; 24: 1019–1033. doi:10.1093/plankt/24.10.1019

66. Kujawinski EB, Longnecker K, Alexander H, Dyhrman ST, Fiore CL, Haley ST, et al. Phosphorus availability regulates intracellular nucleotides in marine eukaryotic phytoplankton. Limnol Oceanogr Lett. 2017; 2: 119–129. doi:10.1002/lol2.10043

67. Van Mooy BAS, Rocap G, Fredricks HF, Evans CT, Devol AH. Sulfolipids dramatically decrease phosphorus demand by picocyanobacteria in oligotrophic marine environments. Proc Natl Acad Sci USA. 2006;103: 8607–8612. doi:10.1073/pnas.0600540103

68. Wurch LL, Bertrand EM, Saito MA, Van Mooy BAS, Dyhrman ST. Proteome Changes Driven by Phosphorus Deficiency and Recovery in the Brown Tide-Forming Alga *Aureococcus anophagefferens*. PLoS One. 2011;6: e28949. doi:10.1371/journal.pone.0028949

69. Peacock MB, Kudela RM. Alkaline phosphatase activity detected in distinct phytoplankton communities in the northern Gulf of Alaska. Mar Ecol Prog Ser. 2013; 473: 79–90. doi:10.2307/24891391

70. Rengefors K, Pettersson K, Blenckner T, Anderson DM. Species-specific alkaline phosphatase activity in freshwater spring phytoplankton: Application of a novel method. J Plankton Res. 2001; 23: 435–443. doi:10.1093/plankt/23.4.435

71. Rengefors K, Ruttenberg KC, Haupert CL, Taylor C, Howes BL, Anderson DM. Experimental investigation of taxon-specific response of alkaline phosphatase activity in natural freshwater phytoplankton. Limnol Ocean. 200; 48(3): 1167–1175

72. Martínez-Soto MC, Basterretxea G, Garcés E, Anglès S, Jordi A, Tovar-Sánchez A. Species-specific variation in the phosphorus nutritional sources by microphytoplankton in a Mediterranean estuary. Front Mar Sci. 2015;54(2). doi:10.3389/fmars.2015.00054

73. Feingersch R, Philosof A, Mejuch T, Glaser F, Alalouf O, Shoham Y, et al. Potential for phosphite and phosphonate utilization by *Prochlorococcus*. ISME J. 2012; 6: 827–834. doi:10.1038/ismej.2011.149

74. Ilikchyan IN, McKay RML, Kutovaya OA, Condon R, Bullerjahn GS. Seasonal Expression of the Picocyanobacterial Phosphonate Transporter Gene phnD in the Sargasso Sea. Front Microbiol. 2010; 135(1). doi:10.3389/fmicb.2010.00135

75. Ilikchyan IN, McKay RML, Zehr JP, Dyhrman ST, Bullerjahn GS. Detection and expression of the phosphonate transporter gene phnD in marine and freshwater picocyanobacteria. Environ Microbiol. 2009; 11: 1314–1324. doi:10.1111/j.1462-2920.2009.01869.x

76. Martínez A, Osburne MS, Sharma AK, Delong EF, Chisholm SW. Phosphite utilization by the marine picocyanobacterium *Prochlorococcus* MIT9301. 2012; 14(6):1363–77. doi:10.1111/j.1462-2920.2011.02612.x

77. Sommer U. The paradox of the plankton: Fluctuations of phosphorus availability maintain diversity of phytoplankton in flow-through cultures1. Limnol Oceanogr. 1984;29: 633–636. doi:10.4319/lo.1984.29.3.0633

78. Droop MR. Some thoughts on nutrient limitation in algae. J Phycol. 1973; 9: 264–272.

79. Kornberg A, Rao NN, Ault-Riché D. Inorganic polyphosphate: a molecule of many functions. Progress in molecular and subcellular biology. 1999; 68: 89–125. doi:10.1146/annurev.biochem.68.1.89

80. Reynolds C. Nutrient uptake and assimilation in phytoplankton. In Ecology of Phytoplankton (Ecology, Biodiversity and Conservation). Cambridge University Press; 2006. p145–177. doi.org/10.1017/CBO9780511542145

81. Ploug H, Stolte W, Jørgensen BB. Diffusive boundary layers of the colony-forming plankton alga Phaeocystis sp.-implications for nutrient uptake and cellular growth. Limnol Oceanogr. 1999; 44: 1959–1967. doi:10.4319/lo.1999.44.8.1959

82. Finkel Z V, Beardall J, Flynn KJ, Quigg A, Alwyn T, Rees V, et al. Phytoplankton in a changing world: cell size and elemental stoichiometry. J plank res. 2010. 32(1):119–137. doi:10.1093/plankt/fbp098

83. Van Veen HW. Phosphate transport in prokaryotes: Molecules, mediators and mechanisms. Antonie van Leeuwenhoek, IJMM. 1997; 72:299–315. doi:10.1023/A:1000530927928

84. Cáceres C, Spatharis S, Kaiserli E, Smeti E, Flowers H, Bonachela JA. Temporal phosphate gradients reveal diverse acclimation responses in phytoplankton phosphate uptake. ISME J. 2019; 13: 2834–2845. doi:10.1038/s41396-019-0473-1

85. Martiny AC, Coleman ML, Chisholm SW. Phosphate acquisition genes in Prochlorococcus ecotypes: Evidence for genome-wide adaptation. Proc Natl Acad Sci U S A. 2006; 103: 12552–12557. doi:10.1073/pnas.0601301103

86. Scanlan DJ, Ostrowski M, Mazard S, Dufresne A, Garczarek L, Hess WR, et al. Ecological Genomics of Marine Picocyanobacteria. Microbiol Mol Biol Rev. 2009; 73: 249–299. doi:10.1128/MMBR.00035-08

87. Chung C-C, Hwang S-PL, Chang J. Identification of a high-affinity phosphate transporter gene in a prasinophyte alga, *Tetraselmis chui*, and its expression under nutrient limitation. Appl Environ Microbiol. 2003; 69: 754–9. doi:10.1128/aem.69.2.754-759.2003

88. Lomas MW, Bonachela JA, Levin SA, Martiny AC. Impact of ocean phytoplankton diversity on phosphate uptake. [cited 3 Apr 2020]. doi:10.1073/pnas.1420760111

89. Grant SR. Phosphorus uptake kinetics and growth of marine osmotrophs. PhD thesis. University of Hawaii at Manoa. 2014.

90. Duhamel S, Moutin T, Van Wambeke F, Van Mooy B, Rimmelin P, Raimbault P, et al. Growth and specific P-uptake rates of bacterial and phytoplanktonic communities in the Southeast Pacific (BIOSOPE cruise). Biogeosciences. 2007; 4: 941–956. doi.org/10.5194/bg-4-941-2007

91. Chisholm SW, Stross RG. Phosphate uptake kinetics in *Euglena gracilis* (Z) (Euglenophyceae) grown on light/dark cycles in synchronised batch cultures. J Phycol. 1976; 12: 210–217. doi:10.1111/j.1529-8817.1976.tb00504.x

92. Rivkin RB, Swift E. Phosphorus metabolism of oceanic dinoflagellates: phosphate uptake, chemical composition and growth of *Pyrocystis noctiluca*. Mar Biol. 1985; 88: 189–198. doi:10.1007/BF00397166

93. Huot Y, Babin M, Bruyant F, Grob C, Twardowski MS, Claustre H. Does chlorophyll a provide the best index of phytoplankton biomass for primary productivity studies? Biogeosciences Discuss. 2007; 4: 707–745. doi:10.5194/bgd-4-707-2007

94. Carstensen A, Herdean A, Schmidt SB, Sharma A, Spetea C, Pribil M, et al. The Impacts of Phosphorus Deficiency on the Photosynthetic Electron Transport Chain. Plant Physiol. 2018; 177: 271–284. doi:10.1104/pp.17.01624

95. Hammond J, PJ White. Sucrose transport in the phloem: integrating root responses to phosphorus starvation. J Exp Bot. 2008; 59: 93–109.

96. Tett P, Cottrell JC, Trew DO, Wood BJB. Phosphorus quota and the chlorophyll: carbon ratio in marine phytoplankton. Limnol Oceanogr. 1975; 20: 587–603. doi:10.4319/lo.1975.20.4.0587

97. Escoubas JM, Lomas M, LaRoche J, Falkowski PG. Light intensity regulation of cab gene transcription is signalled by the redox state of the plastoquinone pool. Proc Natl Acad Sci U S A. 1995; 92: 10237–10241. doi:10.1073/pnas.92.22.10237

98. Geider R, La Roche J. Redfield revisited: variability of C:N:P in marine microalgae and its biochemical basis. Eur J Phycol. 2002; 37: 1–17. doi:10.1017/S0967026201003456

99. Geider R, Graziano L, Mckay & RM. Responses of the photosynthetic apparatus of *Dunaliella tertiolecta* (Chlorophyceae) to nitrogen and phosphorus limitation. Eur J Phycol. 2002; 37(1): 315–332. doi:10.1080/09670269810001736813

100. La Roche J, Geider RJ, Graziano LM, Murray H, Lewis K. Induction of specific proteins in eukaryotic algae grown under Iron-Phosphorus-, or nitrogen deficient conditions. J Phycol. 1993; 29: 767–777. doi:10.1111/j.0022-3646.1993.00767.x

101. Otero-Ferrer JL, Cermeño P, Bode A, Fernández-Castro B, Gasol JM, Morán XAG, et al. Factors controlling the community structure of picoplankton in contrasting marine environments. Biogeosciences. 2018; 15: 6199–6220. doi:10.5194/bg-15-6199-2018

102. Post AF. Nutrient Limitation of Marine Cyanobacteria In: Huisman J., Matthijs H.C., Visser P.M. Editors. Harmful Cyanobacteria. Aquatic Ecology Series, vol 3. Springer, Dordrecht. p 87–107. doi:10.1007/1-4020-3022-3_5

103. Calvo-Díaz A, Morán XAG. Seasonal dynamics of picoplankton in shelf waters of the southern Bay of Biscay. Aquat Microb Ecol. 2006; 42: 159–174. doi:10.3354/ame042159

104. Mousing EA, Richardson K, Ellegaard M. Global patterns in phytoplankton biomass and community size structure in relation to macronutrients in the open ocean. Limnol Oceanogr. 2018; 63: 1298–1312. doi:10.1002/lno.10772

105. Not F, Siano R, Kooistra WHCF, Simon N, Vaulot D, Probert I. Diversity and Ecology of Eukaryotic Marine Phytoplankton. Advances in Botanical Research. Academic Press. 2012; 64: 1–53. doi:10.1016/B978-0-12-391499-6.00001-3

106. Rii YM, Duhamel S, Bidigare RR, Karl DM, Repeta DJ, Church MJ. Diversity and productivity of photosynthetic picoeukaryotes in biogeochemically distinct regions of the South East Pacific Ocean. Limnol Oceanogr. 2016; 61: 806–824. doi:10.1002/lno.10255

107. Righetti D, Vogt M, Gruber N, Psomas A, Zimmermann NE. Global pattern of phytoplankton diversity driven by temperature and environmental variability. Sci Adv. 2019;5(5): eaau6253. doi:10.1126/sciadv.aau6253

108. Brocks JJ, Love GD, Summons RE, Knoll AH, Logan GA, Bowden SA. Biomarker evidence for green and purple sulphur bacteria in a stratified Palaeoproterozoic sea. Nature. 2005; 437: 866–870. doi:10.1038/nature04068

109. Blumenberg M, Thiel V, Riegel W, Kah LC, Reitner J. Biomarkers of black shales formed by microbial mats, Late Mesoproterozoic (1.1Ga) Taoudeni Basin, Mauritania. Precambrian Res. 2012;196–197: 113–127. doi:10.1016/j.precamres.2011.11.010

110. Flannery EN, George SC. Assessing the syngeneity and indigeneity of hydrocarbons in the ~1.4 Ga Velkerri Formation, McArthur Basin, using slice experiments. Org Geochem. 2014; 77: 115–125. doi:10.1016/j.orggeochem.2014.10.008

111. Luo G, Hallmann C, Xie S, Ruan X, Summons RE. Comparative microbial diversity and redox environments of black shale and stromatolite facies in the Mesoproterozoic Xiamaling Formation. Geochim Cosmochim Acta. 2015; 151: 150–167. doi:10.1016/j.gca.2014.12.022

112. Luo Q, George SC, Xu Y, Zhong N. Organic geochemical characteristics of the Mesoproterozoic Hongshuizhuang Formation from northern China: Implications for thermal maturity and biological sources. Org Geochem. 2016; 99: 23–37. doi:10.1016/j.orggeochem.2016.05.004

113. Suslova EA, Parfenova TM, Saraev S V., Nagovitsyn KE. Organic geochemistry of rocks of the Mesoproterozoic Malgin Formation and their depositional environments (southeastern Siberian Platform).Russ Geol Geophys. 2017; 58: 516–528. doi:10.1016/j.rgg.2016.09.027

114. Javaux EJ. Patterns of diversification in early eukaryotes [Modes de diversification des premiers Eucaryotes]. In: Steepman P, Javaux E. editors, Recent Advances in Palynology.-Carnets de Géologie / Notebooks on Geology, Brest, Memoir 2007/01, Abstract 06 (CG2007_M01/06).

115. Leiming Y, Xunlai Y, Fanwei M, Jie H. Protists of the upper mesoproterozoic Ruyang Group in Shanxi Province, China. Precambrian Res. 2005;141: 49–66. doi:10.1016/j.precamres.2005.08.001

116. Moczydłowska M, Landing E, Zang W, Pakacios T. Proterozoic phytoplankton and timing of Chlorophyte algae origins. Palaeontology. 2011; 54: 721–733. doi:10.1111/j.1475-4983.2011.01054.x

117. Agić H, Moczydłowska M, Yin L. Diversity of organic-walled microfossils from the early Mesoproterozoic Ruyang Group, North China Craton –A window into the early eukaryote evolution. Precambrian Res. 2017; 297: 101–130. doi:10.1016/j.precamres.2017.04.042

118. Javaux EJ, Knoll AH. Micropaleontology of the lower Mesoproterozoic Roper Group, Australia, and implications for early eukaryotic evolution. J Paleontol. 2017; 91: 199–229. doi:10.1017/jpa.2016.124

119. Cohen PA, Riedman LA. It’s a protist-eat-protist world: recalcitrance, predation, and evolution in the Tonian–Cryogenian ocean. Emerg Top Life Sci. 2018; 1–8. doi:10.1042/ETLS20170145

120. Fenchel T. Suspension feeding in ciliated protozoa: Functional response and particle size selection. Microb Ecol. 1980; 6: 1–11. doi:10.1007/BF02020370

121. Canter HM, Heaney SI, Lund JWG. The ecological significance of grazing on planktonic populations of cyanobacteria by the ciliate *Nassula*. New Phytol. 1990;114: 247–263. doi:10.1111/j.1469-8137.1990.tb00397.x

122. Sigee DC, Glenn R, Andrews MJ, Bellinger EG, Butler RD, Epton HAS, et al. Biological control of cyanobacteria: Principles and possibilities. Hydrobiologia. 1999; 398:161–172. doi:10.1007/978-94-017-3282-6_15

123. Duhamel S, Björkman KM, Repeta DJ, Karl DM. Phosphorus dynamics in biogeochemically distinct regions of the southeast subtropical Pacific Ocean. Prog Oceanogr. 2017;151: 261–274. doi:10.1016/j.pocean.2016.12.007

